# A Novel Role of Hyaluronan and its Membrane Receptors, CD44 and RHAMM in Obesity-Related Glomerulopathy

**DOI:** 10.1101/2024.06.22.600183

**Authors:** Bingxue Qi, Vishal Musale, Xiong Weng, Ayman K. Banah, Colin E. Murdoch, Abigail C. Lay, Kate J. Heesom, Wenjun Ju, Markus Bitzer, Claire Hills, Richard J.M. Coward, Li Kang

## Abstract

Obesity-related glomerulopathy (ORG) contributes to diabetic nephropathy and kidney cancer, leading to chronic/end-stage kidney disease. To date, treatments for ORG are limited because of incomplete understanding of the disease pathogenesis. Here, we identified a novel role for hyaluronan (HA) and its membrane receptors, CD44 and RHAMM in obesity-associated renal inflammation, fibrosis, tubular injury, and kidney dysfunction. Pharmacological and genetic ablation of HA, CD44 or RHAMM reversed these renal disorders induced by high fat diet feeding in mice *in vivo*. Increased HA content, and CD44 and RHAMM expression damaged the kidney via activation of TGF-β1/Smad2/3, P38/JNK MAPK and ROCK/ERK pathways. We further established a link between renal insulin resistance and ECM remodelling using human kidney cells *in vitro*, shedding mechanistic insight into the role of HA, CD44 and RHAMM in the pathogenesis of ORG. Furthermore, in human kidney biopsies gene expression of CD44 and RHAMM was increased in chronic kidney disease and diabetic nephropathy, and their levels were correlated with markers of kidney (dys)function (GFR, serum creatinine, proteinuria). Our findings provide evidence for HA-CD44/RHAMM as a potential therapeutic target in ORG and consequent prevention of chronic kidney disease.

## Introduction

The incidence of obesity-related glomerulopathy (ORG) is increasing at an alarming rate worldwide in conjunction with the epidemic of obesity. ORG is associated with poor health outcomes including chronic kidney disease (CKD), end-stage renal disease, and increased mortality. Current treatments for ORG (e. g. weight loss and RAAS inhibitors) are limited and attenuate over time because of incomplete understanding of the disease pathogenesis. Therefore, understanding the mechanisms of ORG and identifying new early intervention strategies would have major clinical and socioeconomic benefits.

Extracellular matrix (ECM) remodelling is implicated in the pathogenesis of several metabolic disorders such as cancer[1], type 2 diabetes[2], atherosclerosis[3], obesity[4] and insulin resistance[5]. Hyaluronan (HA), a non-sulfated glycosaminoglycan formed linearly by repeating units of glucuronic acid and acetylglucosamine and one of the major ECM constituents, participates in tissue repair and disease progress[6]. In the healthy kidney, it is mainly found in the interstitium of the inner medulla, where it regulates fluid balance, with limited presence in the outer zone of the outer medulla and virtually void of HA in the cortex[7]. During pathophysiological conditions of kidney-related diseases such as acute kidney injury (AKI)[8], diabetic nephropathy[9], lupus nephritis[10], chronic cyclosporine nephropathy[11] and IgA nephropathy[12], HA accumulation has been associated with renal inflammation and interstitial fibrosis.

CD44 is a major cell surface receptor for HA. The HA-CD44 interaction is important in regulating cell proliferation, aggregation, adhesion, angiogenesis and immune cell migration and activation[13, 14]. Moreover, recent evidence suggests a pivotal role of CD44 in metabolic regulation[15], such as skeletal muscle insulin resistance[5] and obesity[16]. In the kidney, CD44 is mostly localized on the basolateral membranes of collecting ducts in the inner stripes of outer medulla, the thin descending limb of Henle’s loop and macula densa cells[17]. CD44 expression in kidney tissue was positively correlated with renal injury, inflammation, and fibrosis in mice[18]. Genetic deletion of CD44 reduces the number of glomerular lesions in crescentic glomerulonephritis[19]. Oligosaccharides ameliorated AKI by alleviating CD44-mediated immune responses and inflammation in renal tubular cells[20]. The interaction between activated memory T cells and HA/CD44 in the glomerular endothelial glycocalyx is critically important in the glomerular homing of T cells in mouse models of lupus nephritis[21], therefore HA/CD44 plays an important role in mediating pathogenic mechanisms in lupus nephritis[10]. Upregulation of HA/CD44 contributes to interstitial fibrosis in chronic cyclosporin A (CsA)-induced renal injury and dysfunction[11]. Upon HA binding, CD44 interacts with different proteins and recruits intracellular signalling molecules to activate NF-κB,[22] PKC[23], PI3K/Akt[24], Src-ERK[25] and MAPK-signalling pathways[26], subsequently inducing inflammatory processes and regulating cell adhesion, proliferation, migration, and tumour cell invasion.

Receptor for HA mediated motility (RHAMM) is another important HA receptor described by Hardwick et al[27]. RHAMM is poorly expressed in most homeostatic adult tissues. However, RHAMM mRNA and protein expression are strongly, but transiently, increased in response to injury[28]. RHAMM differs from CD44 in structure, but they seem to perform similar or synergistic functions in many diseases[29]. Research indicates that CD44-mediated cellular processes including inflammation, wound healing, tumour formation, and cell migration may require RHAMM surface expression[30]. In contrast to the many studies on CD44 and kidney diseases[19, 20], there are only a few studies on RHAMM, related to renal cell carcinomas[31]. RHAMM participates in inflammation, angiogenesis[32, 33] and tissue response to injury[34], all of which are important factors in the pathogenesis of kidney disease.

We have previously showed that HA accumulation contributed to obesity-associated skeletal muscle insulin resistance, and that reduction of HA or genetic deletion of CD44 improved muscle insulin resistance in obese mice[5, 35]. However, the role of HA and CD44 in regulating kidney function in the setting of obesity or ORG has not been studied. Moreover, the involvement of RHAMM in this process is unknown. In this study, we utilised both genetic and pharmacological approaches to reduce the HA-CD44/RHAMM pathway in mice for renal metabolic and functional phenotyping, together with human kidney cell lines and patient biopsies. Our results reveal a novel role of the HA-CD44/RHAMM pathway in the pathogenesis of ORG, which has the potential for developing new treatment strategies for ORG and its associated kidney diseases.

## Results

### PEGPH20 treatment attenuated obesity-induced renal HA accumulation, tubular damage, renal dysfunction and fibrosis

We have previously shown that in diet-induced obese mice that administering PEGPH20, human recombinant PEGylated hyaluronidase PH-20, at 1mg/kg body weight reduced muscle HA to levels seen in chow-fed mice, decreased fat mass and improved muscle insulin resistance[35]. Here we assessed its role in ORG. We discovered that kidney cortex HA (Fig. 1a and 1b) and outer medulla HA (Fig. 1a and 1c) were significantly increased in the HF diet group and these increments were partially rescued by PEGPH20 treatment. Furthermore, PAS staining showed a significant increase of the glomerular area in the HF diet-fed mice relative to chow-fed controls. However, the glomerular areas were not different between vehicle group and PEGPH20 group on HF diet (Fig. 1a and 1d). Moreover, the tubular damage, including tubular epithelial cell vacuolar deformation/hypertrophy, tubular dilation, loss of brush border and cell lysis mainly occurred in the cortex and outer stripe of the outer medulla in HF diet mice (Fig. 1a). The tubular injury score of HF diet group was significantly higher than that in the chow-fed mice. PEGPH20 treatment partially prevented the tubular damage progression (Fig. 1e). In addition, serum creatinine concentration, a marker of renal dysfunction was increased by HF diet feeding in mice but decreased by PEGPH20 treatment (Fig. 1f). Renal fibrosis, assessed by collagen deposition and protein expression of α-SMA was increased in HF-fed mice relative to chow-fed controls, which was partially blocked by PEGPH20 treatment (Fig. 1a, 1g and 1h). These results demonstrate that HF diet feeding contributed to renal dysfunction, including HA accumulation, tubular injury, elevated serum creatinine levels and fibrosis, which were partially prevented by PEGPH20 treatment.

**Fig. 1.**
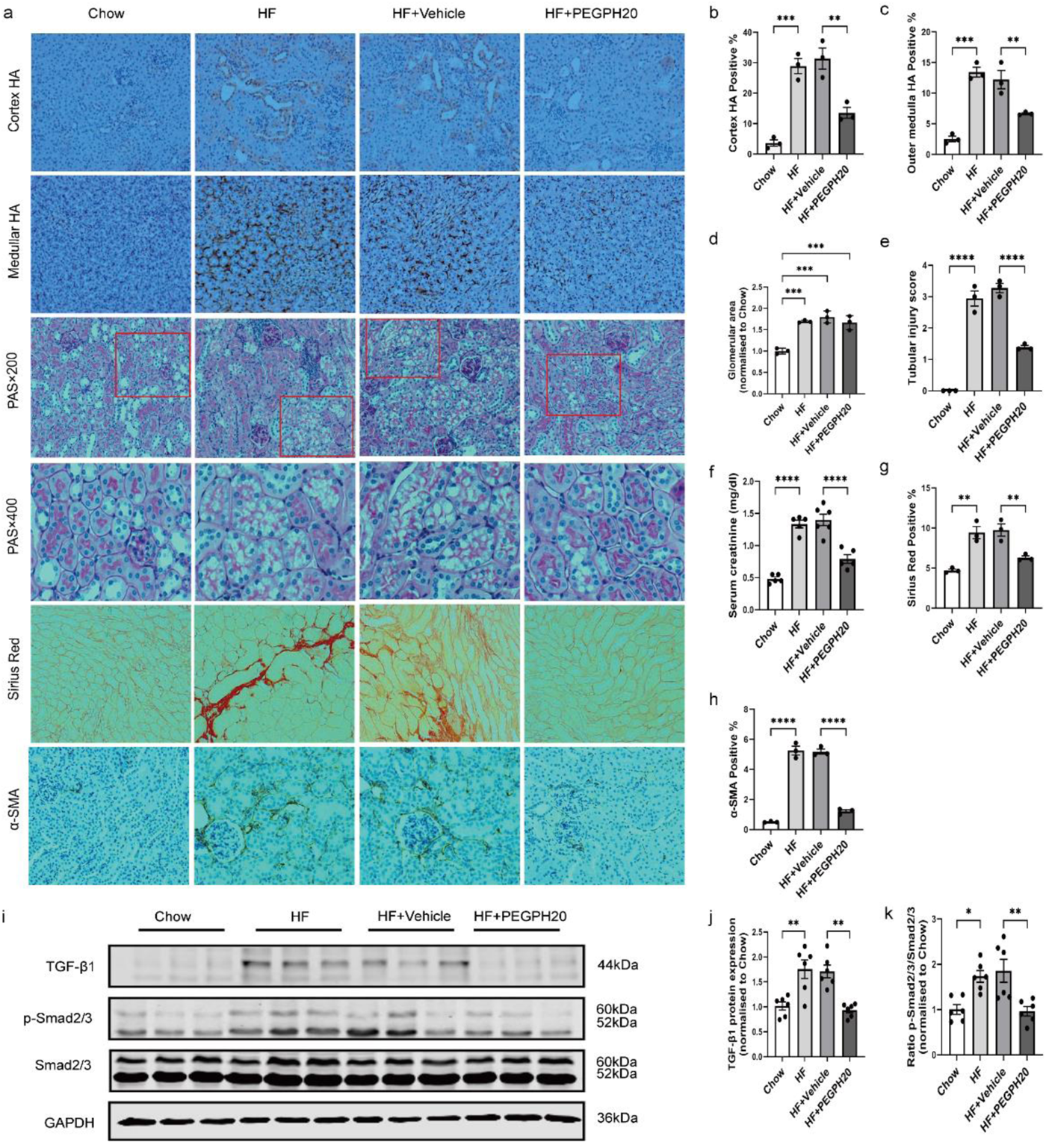
PEGPH20 treatment attenuated obesity-induced HA accumulation, glomerular area expansion, tubular damage, renal dysfunction and fibrosis. **a** Images of HA, PAS, Sirius Red and α-SMA staining in kidneys from chow-fed mice, HF-fed mice and HF-fed mice with either vehicle or PEGPH20 (1mg/kg) treatment. Representative images (×20 magnification) were shown for HA, Sirius Red and α-SMA staining and images (×20 and ×40 magnification) were shown for PAS staining. Areas of HA positive (%), glomeruli, Sirius red positive (%) and α-SMA positive (%) were measured using the Image J program. **b-c** Quantification of HA content. d Glomerular area. **e** Tubular injury score. **f** Serum creatinine concentration. **g** Quantification of collagen deposition. **h** Quantification of α-SMA expression. **i-k** Representative western blot and quantification of protein expression of TGF-β1, p-Smad2/3 and total Smad 2/3 in the kidney. N = 3-6 male mice. Data are mean ± SEM. For all panels, one-way ANOVA followed by Tukey’s post-test for multiple comparisons was used for the analysis of statistical significance. Significance \**p* < 0.05, \*\**p* < 0.01, \*\*\**p* < 0.001, \*\*\*\**p* < 0.0001. Source data are provided in the source data file.

### PEGPH20 blocked the activation of TGF-β1/Smad2/3, P38/JNK MAPK and HA/CD44 pathways and prevented obesity-induced inflammation

We further investigated the mechanisms by which PEGPH20 ameliorated ORG. TGF-β1/Smad2/3 signalling was increased by HF diet feeding in mice, and this was abolished by PEGPH20 treatment (Fig. 1i-k). Moreover, the P38/JNK MAPK phosphorylation levels were increased in the HF diet-fed mice, which was blocked in mice treated with PEGPH20 (Fig. 2a-c). HA is a major component of the renal ECM and it binds to membrane receptor CD44 and RHAMM regulating physiological and pathological activities. We further investigated whether the renal protective effect of PEGPH20 was dependent upon CD44 and RHAMM. HF diet increased CD44 and RHAMM expression and PEGPH20 treatment reduced CD44 expression (Fig. 2a and 2d) in the kidney of HF-fed mice but did not affect RHAMM expression (Fig. 2a and 2e). Furthermore, HF diet feeding resulted in an increase, which was abolished by PEGPH20 in ROCK2 expression and ERK/Akt phosphorylation levels (Fig. 2a and 2f-h), which are important intracellular signalling molecules of the HA-CD44 signalling[15]. Furthermore, mRNA expression of the pro-inflammatory cytokines IL-1β, TNF-α, and IL-6 were increased by HF diet and expression of the anti-inflammatory cytokine IL-10 was decreased by HF diet (Fig 2i-l). These changes were all reversed by PEGPH20 treatment. Collectively, these results suggest that PEGPH20 treatment ameliorated obesity-induced renal injury by suppressing the activation of TGF-β1/Smad2/3, P38/JNK MAPK and HA/CD44 pathways and subsequent inflammation cascade.

**Fig. 2.**
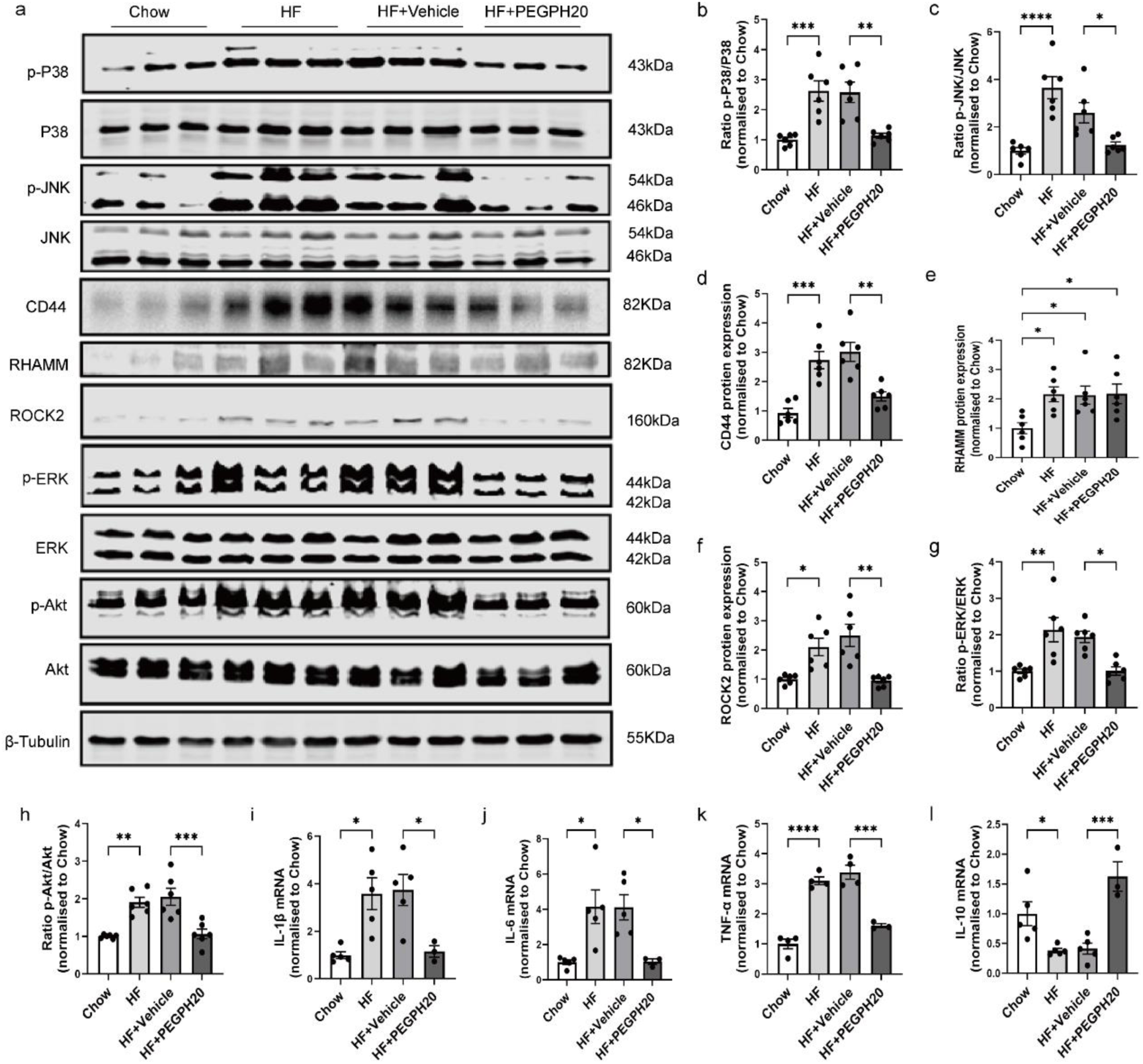
PEGPH20 blocked the activation of the P38/JNK MAPK and HA/CD44 pathways and prevented obesity-induced inflammation. **a-h** Representative western blot images and quantification of protein expression of the P38/JNK MAPK pathway, CD44/RHAMM and HA/CD44 pathway-associated proteins (ROCK2/ERK/Akt) in kidneys from chow-fed mice, HF-fed mice and HF-fed mice with either vehicle or PEGPH20 (1mg/kg) treatment. N = 6 male mice. **i-l** mRNA expression of IL-1β (**i**), IL-6 (**j**), TNF-α (**k**) and IL-10 (**l**) assessed by qRT–PCR. N = 3-5 male mice. Data are mean ± SEM. For all panels, one-way ANOVA followed by Tukey’s post-test for multiple comparisons was used for the analysis of statistical significance. Significance \**p* < 0.05, \*\**p* < 0.01, \*\*\**p* < 0.001, \*\*\*\**p* < 0.0001. Source data are provided in the source data file.

### Global *Cd44* gene deletion attenuated obesity-induced HA accumulation, tubular damage, renal dysfunction and fibrosis

To elucidate whether the CD44 receptor mediated renal injury in ORG, *Cd44*^+/+^ and *Cd44*^-/-^ mice were fed with HF diet for 16 weeks. *Cd44*^-/-^ mice also received either vehicle or PEGPH20 once every 3 days for 28 days. We have previously reported that CD44 contributes to HA-mediated muscle insulin resistance and that the metabolic beneficial effects of PEGPH20 are CD44 dependent[5]. Here, we show that global deletion of *Cd44* gene in HF-fed mice significantly decreased renal cortex and outer medulla HA accumulation (Fig 3a-c). Furthermore, PEGPH20 treatment in *Cd44*^-/-^ mice caused a further reduction in HA accumulation in the outer medulla but not in the cortex. Global deletion of *Cd44* gene in HF-fed mice significantly decreased the tubular injury scores (Fig. 3a and 3e), serum creatinine levels (Fig. 3f), collagen deposition (Fig. 3a and 3g) and α-SMA expression (Fig. 3a and 3h) without affecting the glomerular areas (Fig. 3a and 3d). PEGPH20 treatment caused a further reduction in serum creatinine level in HF-fed *Cd44*^-/-^ mice (Fig. 3f). Moreover, the protein expression of TGF-β1 and phosphorylation of Smad2/3 were reduced in HF-fed *Cd44*^-/-^ mice relative to HF-fed *Cd44*^+/+^ mice and further decreased after PEGPH20 treatment (Fig. 3i-k). These results demonstrate that *Cd44* deletion reduced obesity induced kidney injury, which was potentiated by also reducing HA levels.

**Fig. 3.**
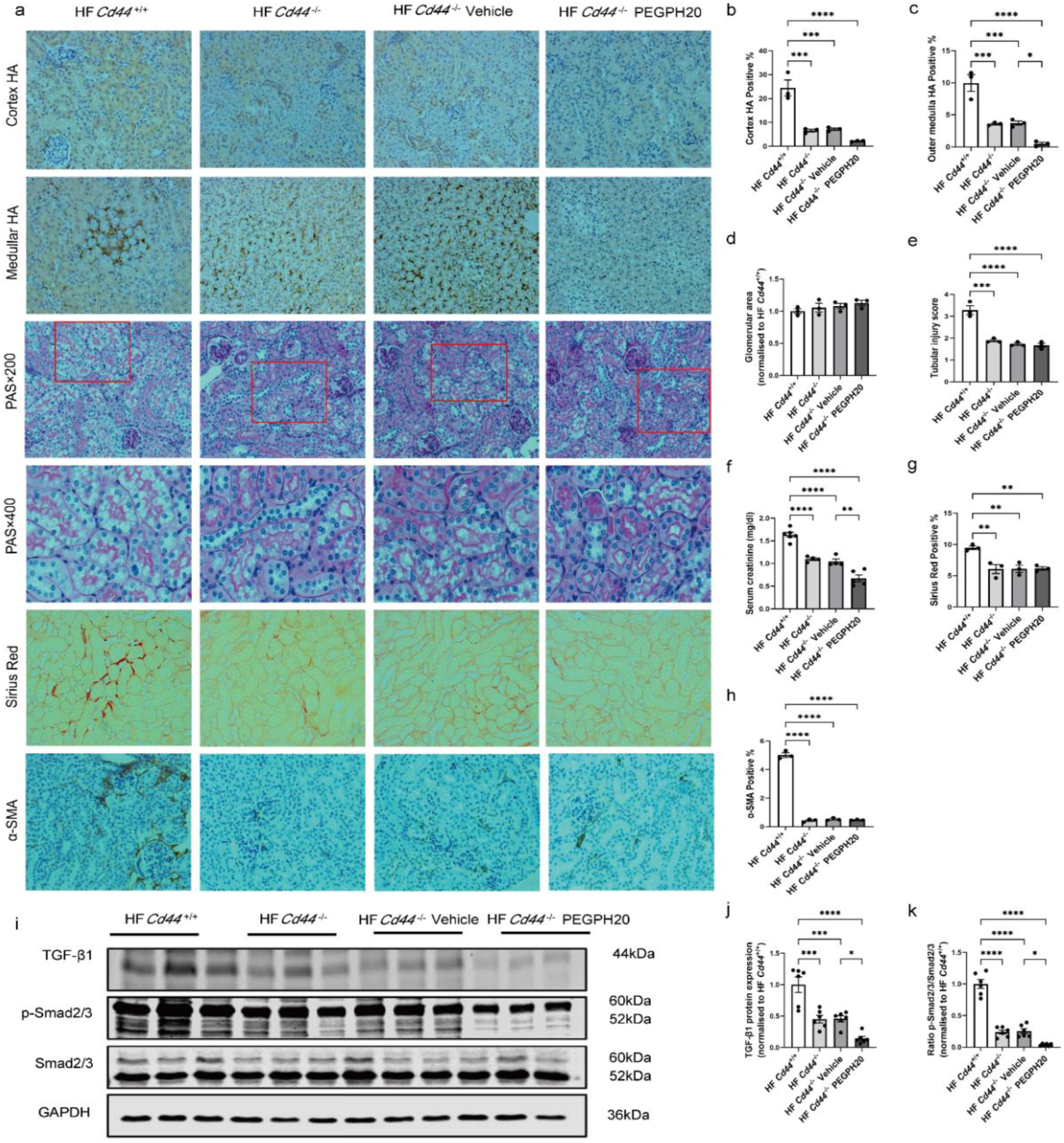
Global *Cd44* gene deletion attenuated obesity-induced HA accumulation, tubular damage, renal dysfunction and fibrosis. **a** Images of HA, PAS, Sirius Red and α-SMA staining in kidneys from HF-fed *Cd44*^+/+^ mice, HF-fed *Cd44*^-/-^ mice and HF-fed *Cd44*^-/-^ mice with either vehicle or PEGPH20 (1mg/kg) treatment. Representative images (×20 magnification) were shown for HA, Sirius Red and α-SMA staining and images (×20 and ×40 magnification) were shown for PAS staining. Areas of HA positive (%), glomeruli, Sirius red positive (%) and α-SMA positive (%) were measured using the Image J program. **b-c** Quantification of HA content. **d** Glomerular area. **e** Tubular injury score. **f** Serum creatinine concentration. **g** Quantification of collagen deposition. **h** Quantification of α-SMA expression. **i-k** Representative western blot images and quantification of protein expression of TGF-β1, p-Smad2/3 and Smad 2/3 in the kidney. N = 3-6 male mice. Data are mean ± SEM. For all panels, one-way ANOVA followed by Tukey’s post-test for multiple comparisons was used for the analysis of statistical significance. Significance \**p* < 0.05, \*\**p* < 0.01, \*\*\**p* < 0.001, \*\*\*\**p* < 0.0001. Source data are provided in the source data file.

Global *Cd44* gene deletion blocked the activation of P38/JNK MAPK and HA/CD44 pathways and prevented obesity-induced inflammation. Global deletion of *Cd44* gene significantly lowered the P38/JNK MAPK phosphorylation levels and this was more pronounced in *Cd44*^-/-^ mice also treated with PEGPH20 (Fig. 4a-c). Furthermore, deletion of the *Cd44* gene significantly reduced the protein expression of CD44 (Fig. 4a and 4d) but had no effect on the protein expression of RHAMM (Fig. 4a and 4e) in the kidney. ROCK2 and ERK/Akt phosphorylation levels were downregulated in HF-fed *Cd44*^-/-^ mice when compared with HF-fed *Cd44*^+/+^ mice, and PEGPH20 had no further effects (Fig. 4a and 4f-h). mRNA levels of IL-1β, TNF-α and IL-6 were markedly decreased, and IL-10 mRNA level was increased in the HF-fed *Cd44*^-/-^ mice in comparison to HF-fed *Cd44*^+/+^ mice. PEGPH20 treatment of *Cd44*^-/-^ mice further reduced IL-1β, TNF-α and IL-6 mRNA levels and increased IL-10 mRNA level compared to vehicle treated *Cd44*^-/-^ mice (Fig. 4i-l). This suggests that CD44 deletion suppresses the activation of TGF-β1/Smad2/3, P38/JNK MAPK and HA/CD44 pathways and subsequent inflammation cascade which attenuates obesity-induced renal injury but this is augmented through HA enzymatic degradation with PEGPH20.

**Fig. 4.**
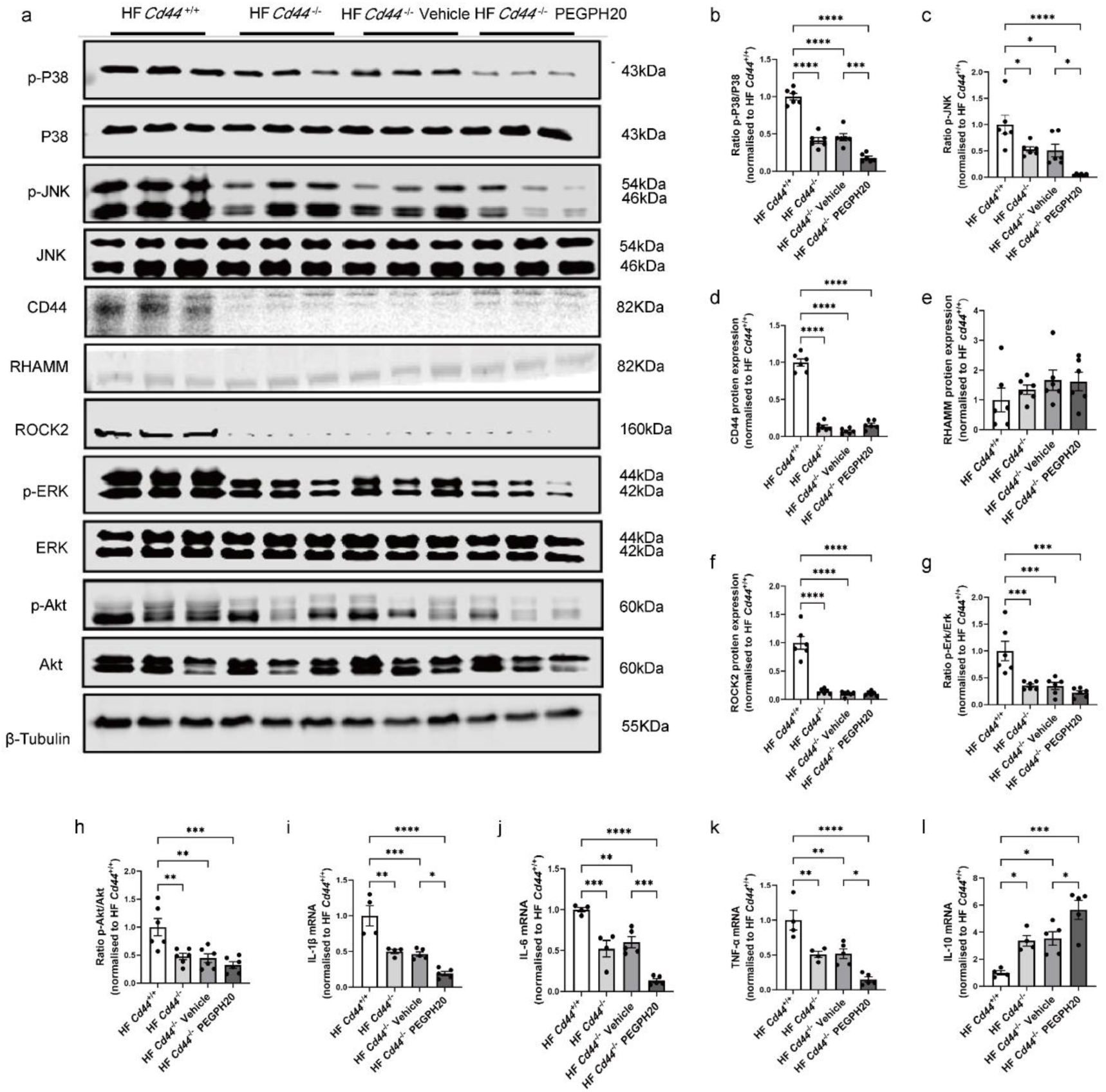
Global *Cd44* gene deletion blocked the activation of P38/JNK MAPK and HA/CD44 pathways and prevented obesity-induced inflammation. **a-h** Representative western blot images and quantification of protein expression of the P38/JNK MAPK pathway, CD44/RHAMM and HA/CD44 pathway-associated proteins (ROCK2/ERK/Akt) in kidneys from HF-fed *Cd44*^+/+^ mice, HF-fed *Cd44*^-/-^ mice and HF-fed *Cd44*^-/-^ mice with either vehicle or PEGPH20 (1mg/kg) treatment. N = 6 male mice. **i-l** mRNA expression of IL-1β (**i**), IL-6 (**j**), TNF-α (**k**) and IL-10 (**l**) assessed by qRT– PCR. N = 3-5 male mice. Data are mean ± SEM. For all panels, one-way ANOVA followed by Tukey’s post-test for multiple comparisons was used for the analysis of statistical significance. Significance \**p* < 0.05, \*\**p* < 0.01, \*\*\**p* < 0.001, \*\*\*\**p* < 0.0001. Source data are provided in the source data file.

### Global *Hmmr* gene deletion reduces obesity-induced HA accumulation, glomerular area expansion, tubular damage, renal dysfunction and fibrosis

The role of RHAMM in renal injury of ORG was explored using *Hmmr*-null mice (*Hmmr*^-/-^) and their wildtype littermate controls (*Hmmr* ^+/+^) fed with an HF or chow diet for 16 weeks. *Hmmr* ^-/-^ and *Hmmr* ^+/+^ mice body weights were not different on either the chow or HF diet (Supplemental Fig. 2a and 2b). Systolic and diastolic blood pressures were lower in chow-fed *Hmmr* ^-/-^ mice compared to chow-fed *Hmmr* ^+/+^ mice, but were similar between genotypes on HF diet (Supplemental Fig. 2c and 2d). In the renal cortex and outer medulla, a similar level of HA was observed in chow-fed *Hmmr* ^+/+^ and *Hmmr* ^-/-^ mice. However, HF diet significantly increased HA accumulation in *Hmmr* ^+/+^ mice, but was significantly reduced in the *Hmmr* ^-/-^ mice (Fig. 5a-c). *Hmmr* gene deletion significantly attenuated HF diet-induced increases in glomerular area, tubular injury score, serum creatinine level, collagen deposition, α-SMA expression, TGF-β1 expression and phosphorylation of Smad2/3 (Fig. 5a and 5d-k).

**Fig. 5.**
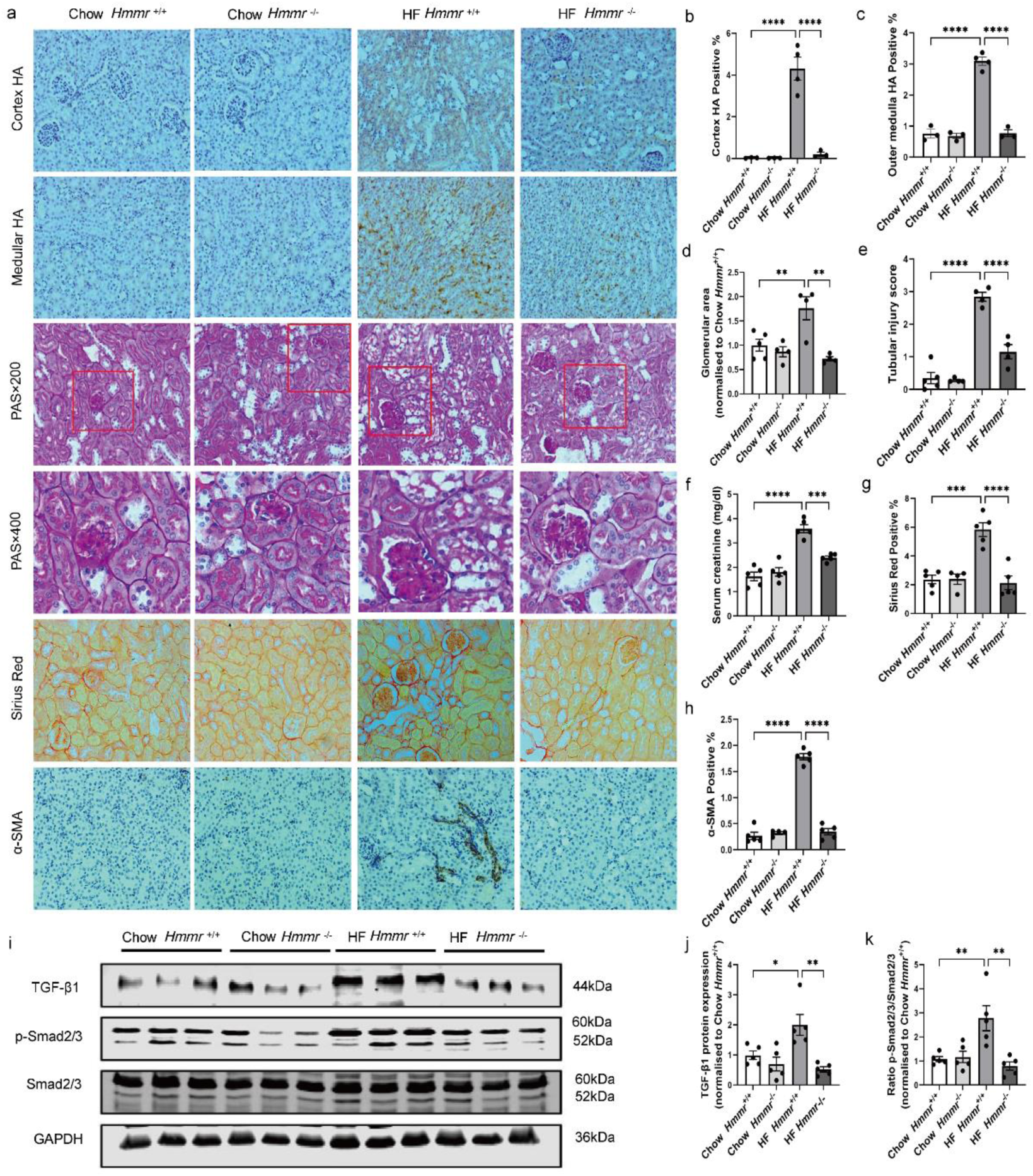
Global *Hmmr* gene deletion attenuated obesity-induced HA accumulation, glomerular area expansion, tubular damage, renal dysfunction and fibrosis. **a** Images of HA, PAS, Sirius Red and α-SMA staining in kidneys from *Hmmr*^+/+^ mice and *Hmmr*^-/-^ mice, both with either chow or high fat diet for 16 weeks. Representative images (×20 magnification) were shown for HA, Sirius Red and α-SMA staining and images (×20 and ×40 magnification) were shown for PAS staining. Areas of HA positive (%), glomeruli, Sirius red positive (%) and α-SMA positive (%) were measured using the Image J program. **b-c** Quantification of HA content. **d** Glomerular area. **e** Tubular injury score. **f** Serum creatinine concentration. **g** Quantification of collagen deposition. **h** Quantification of α-SMA expression. **i-k** Representative western blot images and quantification of protein expression of TGF-β1, p-Smad2/3 and Smad 2/3 in the kidney. N = 3-5 male mice. Data are mean ± SEM. For all panels, one-way ANOVA followed by Tukey’s post-test for multiple comparisons was used for the analysis of statistical significance. Significance \**p* < 0.05, \*\**p* < 0.01, \*\*\**p* < 0.001, \*\*\*\**p* < 0.0001. Source data are provided in the source data file.

### Global *Hmmr* gene deletion blocked the activation of P38/JNK MAPK and HA/CD44 pathways and prevented obesity-induced inflammation

HF-fed *Hmmr*^+/+^ mice significantly increased their renal P38/JNK MAPK phosphorylation (Fig. 6a-c), CD44 and RHAMM protein expression (Fig. 6a,6d, 6e). Additionally, ROCK2 expression (Fig. 6a, 6f) and the levels of both Akt and ERK phosphorylation (Fig. 6a, 6g-h) were significantly increased. These were all significantly reduced in the HF-fed *Hmmr* ^-/-^ mice. The cytokines IL-1β, TNF-α and IL-6 mRNA levels were significantly increased in the HF-*Hmmr*^+/+^ mice whilst IL-10 was decreased. These cytokine alterations were also all reversed in the HF-fed *Hmmr* ^-/-^ mice. These results suggest that *Hmmr* gene deletion alleviates obesity-induced kidney injury possibly by inhibiting the activation of TGF-β1/Smad2/3, P38/JNK MAPK and HA/CD44 pathways and subsequent inflammation. In addition, we speculated that the renal protective effect of *Hmmr* gene deletion was accompanied by reduction of CD44.

**Fig. 6.**
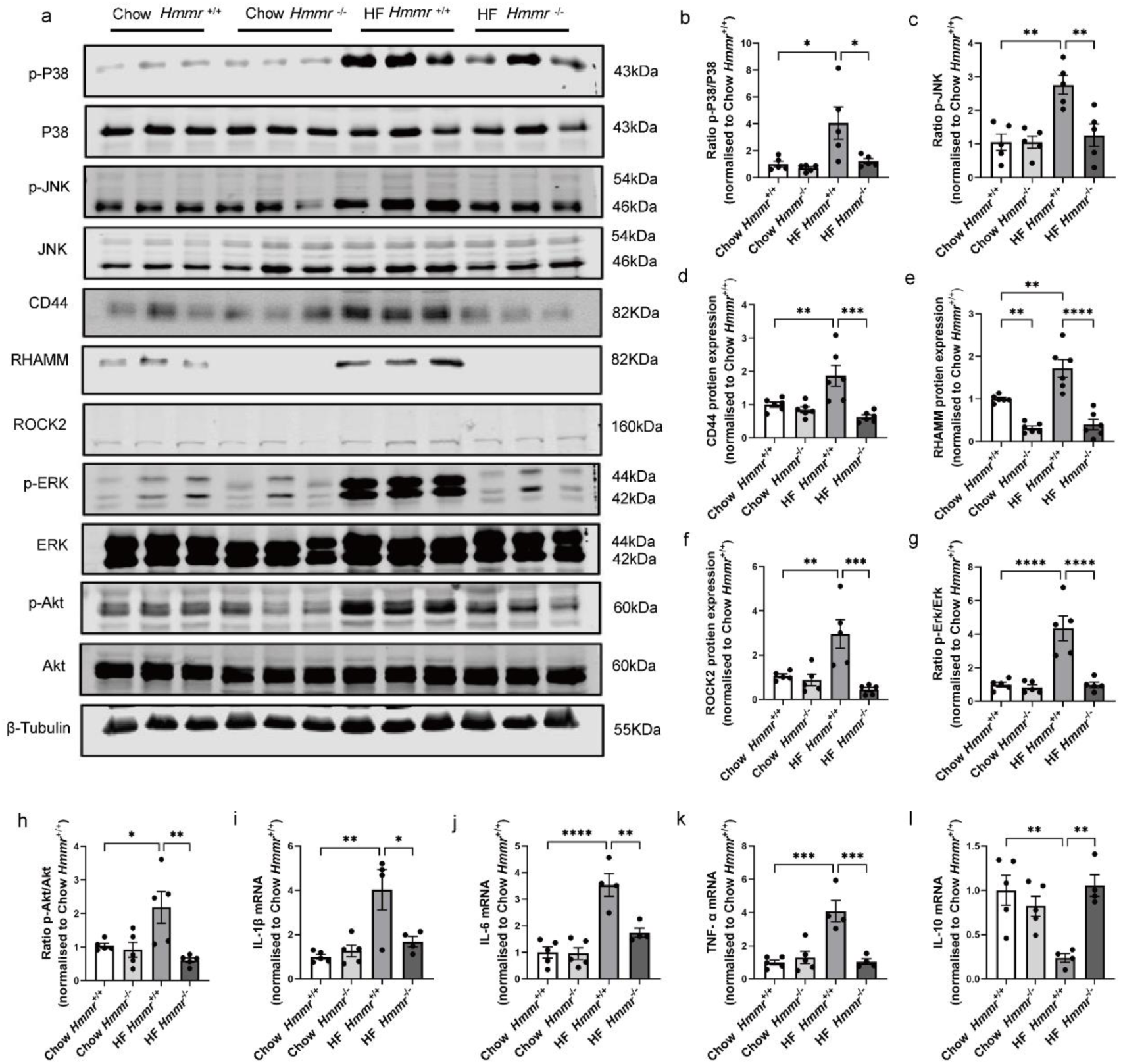
Global *Hmmr* gene deletion blocked the activation of P38/JNK MAPK and HA/CD44 pathways and prevented obesity-induced inflammation. **a-h** Representative western blot images and quantification of protein expression of the P38/JNK MAPK pathway, CD44/RHAMM and HA/CD44 pathway-associated proteins (ROCK2/Erk/Akt) in kidneys from *Hmmr*^+/+^ mice and *Hmmr*^-/-^ mice, both with either chow or high fat diet for 16 weeks. N = 5 male mice. **i-l** mRNA expression of IL-1β (**i**), IL-6 (**j**), TNF-α (**k**) and IL-10 (**l**) assessed by qRT–PCR. N = 4-5 male mice. Data are mean ± SEM. For all panels, one-way ANOVA followed by Tukey’s post-test for multiple comparisons was used for the analysis of statistical significance. Significance \**p* < 0.05, \*\**p* < 0.01, \*\*\**p* < 0.001, \*\*\*\**p* < 0.0001. Source data are provided in the source data file.

### CD44 and RHAMM were upregulated in insulin resistant human kidney cells

We next investigated which cell types in the kidney are responsible for obesity-associated renal injury using conditionally immortalized human kidney cells[36]. Obesogenic and insulin resistant conditions were induced by incubating cells with high glucose, high insulin and inflammatory cytokines (i.e. TNF-α and IL-6)[37]. Proteomic analysis revealed that CD44 protein expression was increased in insulin resistant podocytes, proximal tubular (PT) cells and mesangial cells, but not glomerular endothelial cells (GEnC) (Fig. 7a). Interestingly, RHAMM expression was only increased in insulin resistant podocytes (Fig. 7b). HA remodelling was observed in insulin resistant podocytes and PT cells as intracellular HA binding protein (HABP) was decreased in insulin resistant PT cells whereas cell surface hyaluronidase expression was increased in insulin resistant podocytes (Fig. 7c-d). Moreover, insulin resistance caused a considerable collagen deposition in podocytes, PT and GEnC cells as evidenced by increased expression of various collagen isoforms, but not in the mesangial cells (Fig. 7e-k). Consistently, the protein expression of collagen receptor integrins was also increased by insulin resistance, mostly in PT cells and to a less extend in podocytes and mesangial cells, but not in the GEnC cells (Fig. 7l-p). Intriguingly, collagen remodelling (e.g. glycosylation and crosslinking) was induced by insulin resistance mainly in the GEnC cells as evidenced by increased expression of collagen modifying enzymes (Fig. 7q-t). These results exhibit human relevance of the mouse work and provide insight into the role of different cell types of the kidney in obesity and insulin resistance-associated renal injury.

**Fig. 7.**
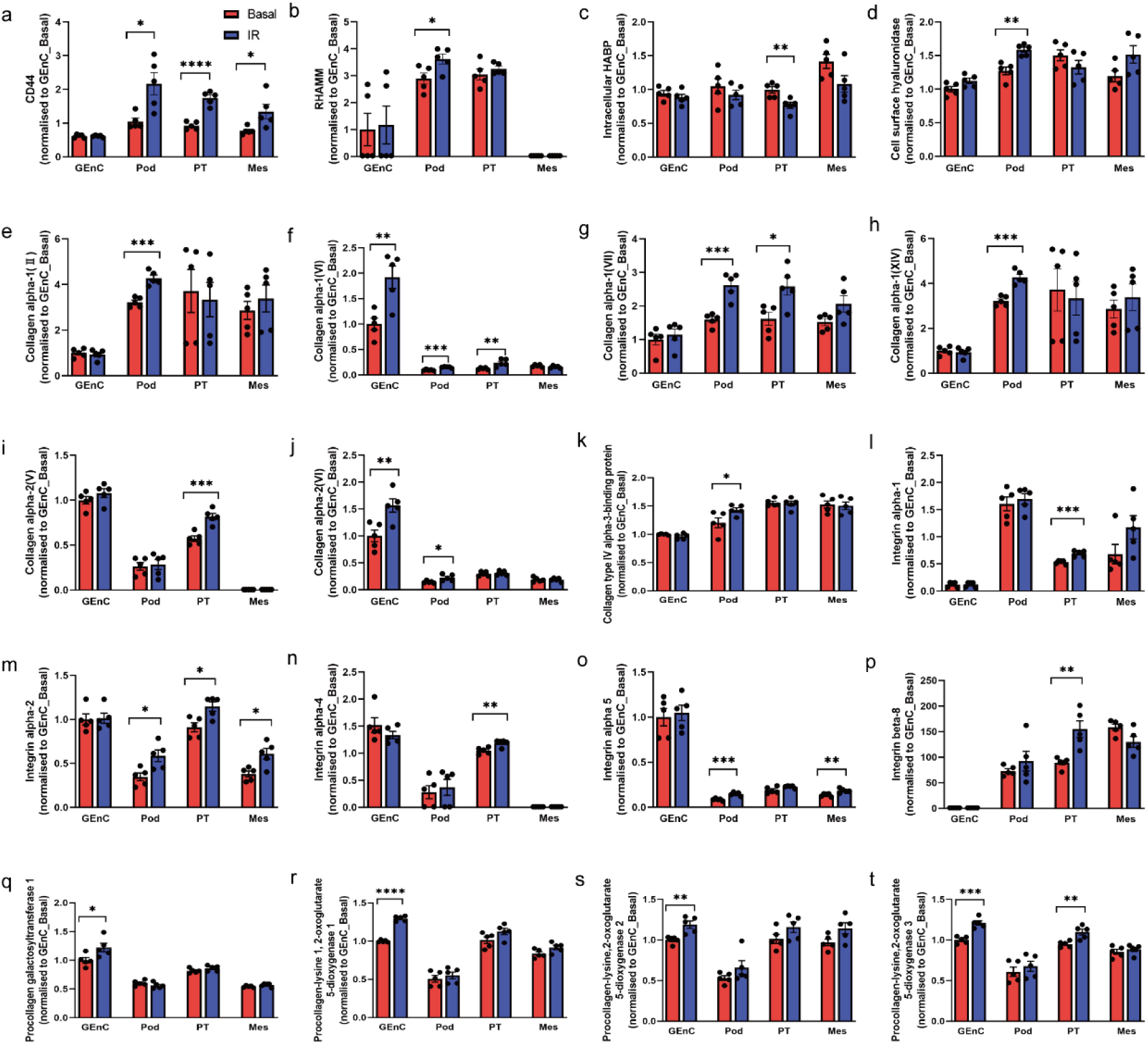
CD44 and RHAMM were upregulated in insulin resistant human kidney cells. Proteomics analysis of protein expression in basal vs insulin resistant glomerular endothelial cells (GEnC), podocytes (Pod), proximal tubular cells (PT), and mesangial cells (MES). **a** CD44; **b** RHAMM; **c** Intracellular HA binding protein (HABP); **d** Cell surface hyaluronidase; **e-k** Protein expression of various collagen isoforms. Collagen alpha-1 (II) (**e**), Collagen alpha-1 (VI) (**f**), Collagen alpha-1 (VII) (**g**), collagen alpha-1 (XIV) (**h**), Collagen alpha-2 (V) (**i**), Collagen alpha-2 (VI) (**j**) and Collagen type IV alpha-3-binging protein (**k**); **l-p** Protein expression of collagen receptor integrins alpha-1 (**l**), integrins alpha-2 (**m**), integrins alpha-4 (**n**), integrins alpha-5 (**o**) and integrins beta-8 (**p**); **q-t** Protein expression of collagen modifying enzymes procollagen galactosyltransferase 1 (**q**), procollagen-lysine 1, 2-oxoglutarate 5-dioxygenase 1 (**r**), procollagen-lysine,2-oxoglutarate 5-dioxygenase 2 (**s**) and procollagen-lysine,2-oxoglutarate 5-dioxygenase 3 (**t**). N = 5. Data are mean ± SEM. For all panels, the unpaired, two-tailed Student’s t test was used for the analysis of statistical significance between basal and insulin resistant cells. Significance \**p* < 0.05, \*\**p* < 0.01, \*\*\**p* < 0.001. Source data are provided in the source data file.

### Increased CD44 and RHAMM expression correlated with a decline in kidney function in humans

Using publically available transcriptomic datasets from the Nephroseq repository, we found that gene expression of CD44 and RHAMM was increased in kidney biopsies of patients with CKD and diabetic nephropathy when compared to those from healthy living donors (Fig. 8a-f). Functionally, CD44 expression was negatively correlated with GFR, but positively correlated with serum creatinine levels in patients with diabetic nephropathy (Fig. 8g-h). Likewise, RHAMM expression was positively correlated with proteinuria in patients with diabetic nephropathy (Fig. 8i).

**Fig 8.**
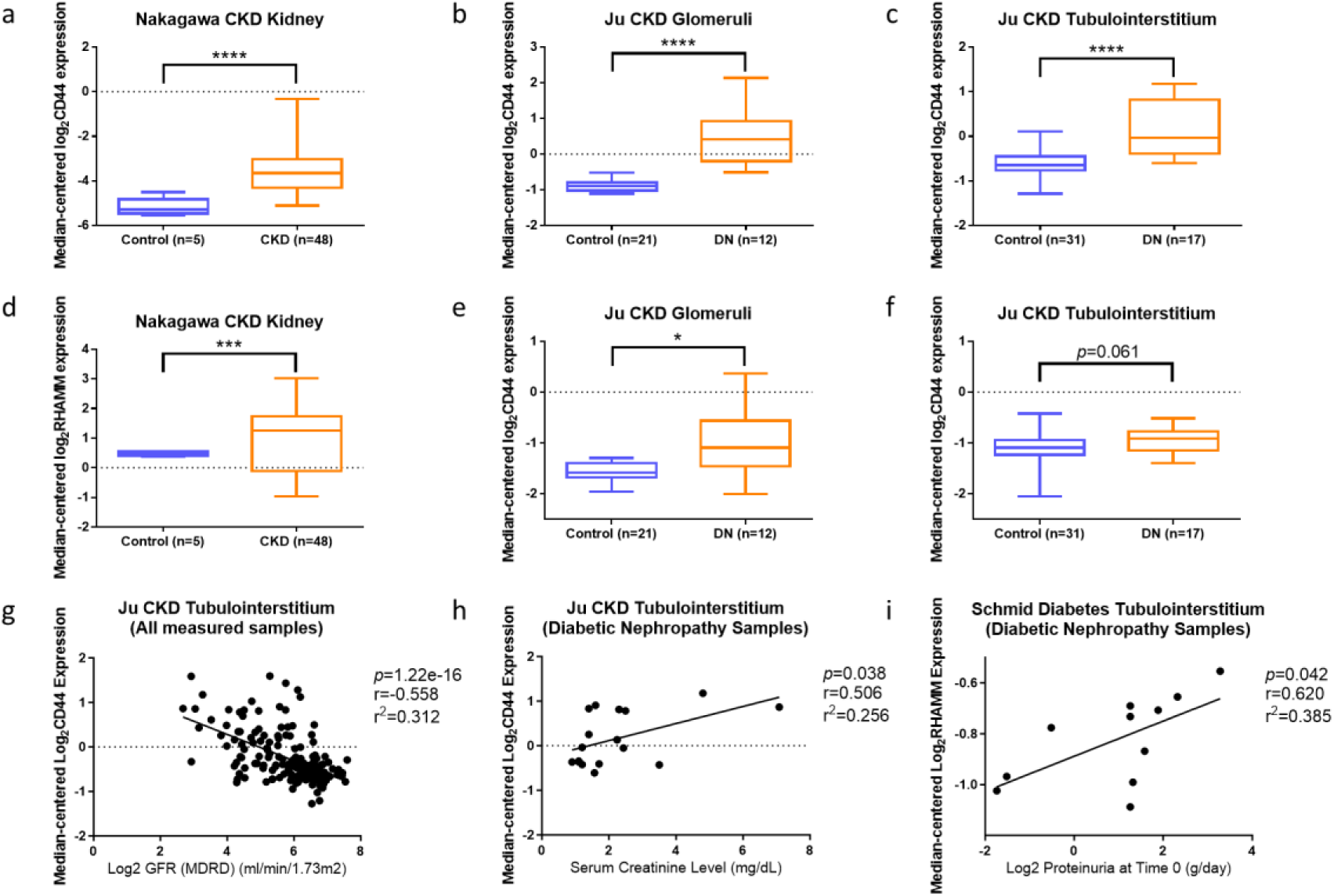
Increased CD44 and RHAMM expression correlated with a decline in kidney function in humans. **a-f** CD44 and RHAMM gene expression was analysed by Nephroseq v5 using datasets of Nakagawa CKD (chronic kidney disease) Kidney[67] and Ju CKD Glomeruli and Tubulointerstitium[68]. Two-tailed student’s t-tests were used for satistical comparision. **g-i** Correlations of CD44 and RHAMM gene expression with GFR, serum creatinine level and proteinuria were analysed by Nephroseq v5 using datasets of Ju CKD Tubulointerstium and Schmid Diabetes Tubulointerstitum[69]. Pearson correlations were performed for statistical analysis with p measuring statistical significance, r measuring the linear correlation between two variables and r^2^ measuring how close the data are to the fitted regression line. Significance \**p* < 0.05, \*\*\**p* < 0.005, \*\*\*\**p* < 0.001.

## Discussion

In the present study, we showed that HF diet feeding in mice led to renal HA accumulation, renal inflammation and fibrosis, and consequent renal injury. Reduction of renal HA by human recombinant hyaluronidase prevented these HF diet-induced renal disorders. Moreover, genetic deletion of CD44 or RHAMM, the two membrane receptors for HA also provided renal protective effects in HF diet-induced obese mice. These results demonstrate a novel role for HA and its receptors in ORG in mice and we further showed that the HA-CD44/RHAMM pathway may cause renal damage by activating the TGF-β1/Smad2/3, P38/JNK MAPK, and ROCK2/ERK/Akt pathways (Fig. 9).

**Fig 9.**
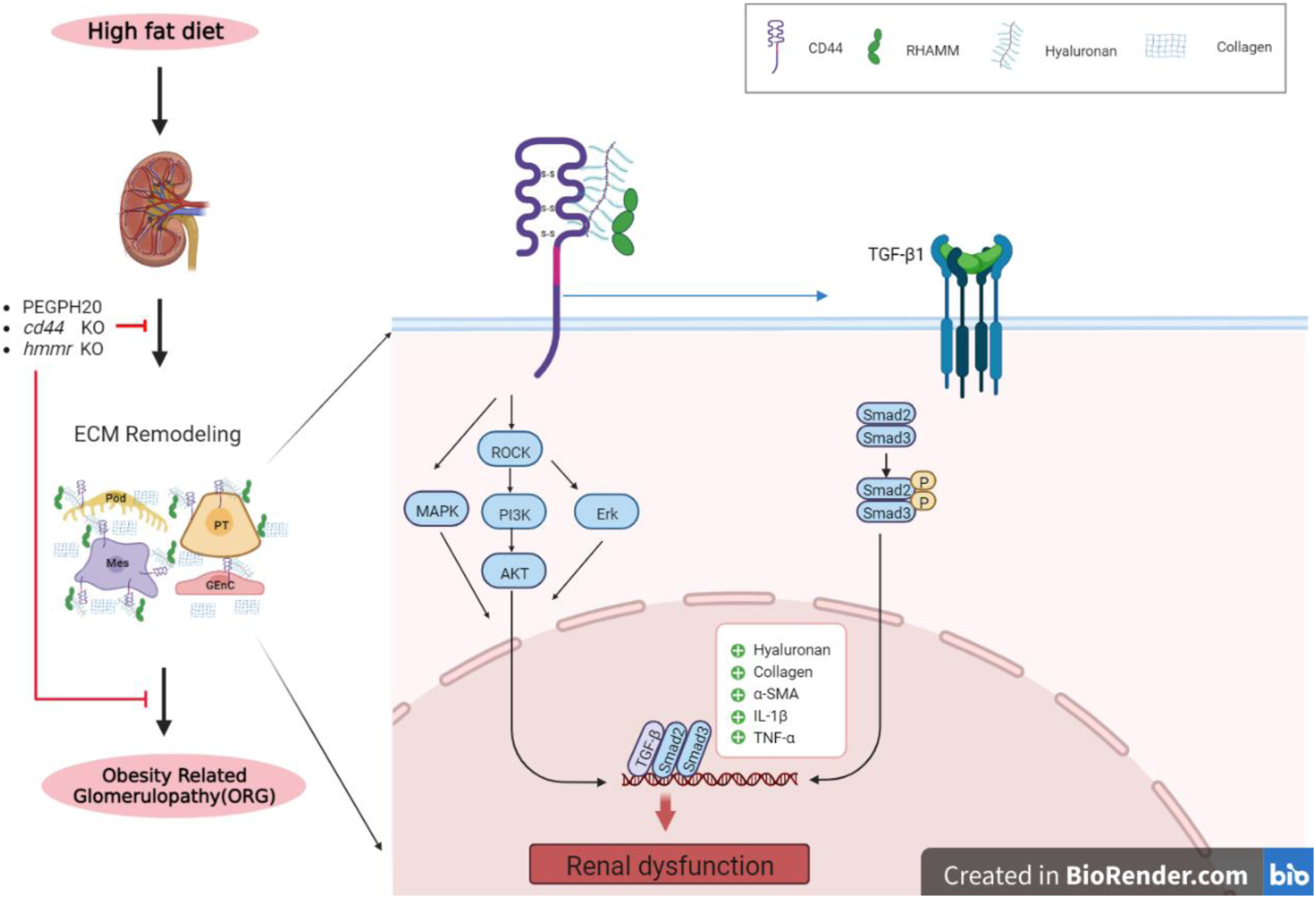
Working model for HA-CD44/RHAMM pathway in the pathogenesis of ORG. Obesity-induced renal ECM remodelling. PEGPH20 treatment and/or global *cd44*/*Hmmr* genes deletion decreased HA accumulation and blocked the activation of TGF-β1/Smad2/3, P38/JNK MAPK and HA/CD44 pathways and subsequent inflammation, prevented obesity-induced inflammation and partially attenuated ECM remodelling induced renal dysfunction.

ORG, a risk factor for the development of CKD is characterised by glomerular hypertrophy, mesangial matrix expansion, and focal segmental glomerulosclerosis[38]. Another cardinal feature of ORG is the accumulation of lipid vacuoles in podocytes, mesangial cells, and PT cells with structural and functional changes, suggesting that abnormal lipid metabolism and lipotoxicity may be the major cause of renal dysfunction and activation of proinflammatory processes[39]. These histologic changes are consistent with what we have observed in the present study. Our study showed a significant increase of the glomerular area in the HF diet-fed mice. Furthermore, the tubular damage, including tubular epithelial cell vacuolar deformation/hypertrophy, tubular dilation, loss of brush border and cell lysis also occurred in mice after 16 weeks of HF diet feeding with an increased serum creatinine concentration. These results demonstrate that HF diet feeding in mice reassembles pathologies of human ORG, therefore representing a valuable experimental model.

Upon kidney injury, profibrotic factors can be secreted by injured tubular epithelia and infiltrated inflammatory cells to promote fibroblast differentiation into myofibroblasts and ECM production, leading to renal fibrosis. Tubulointerstitial fibrosis was observed in obesity-related kidney disease[40]. Renal fibrosis is characterized by pathological deposition of the ECM, which disrupts normal kidney architecture and damages renal function[41]. The ECM HA was shown to be induced in the initial response phase of kidney injury and remodelled during disease progression of AKI, CKD, and diabetic nephropathy[42], suggesting a pathological role of HA in kidney injury. Here we also showed increased HA accumulation in ORG in mice. Fibroblast differentiation into myofibroblasts is regulated by classical TGF-β1-dependent Smad-signalling pathway, yet TGF-β1-mediated fibroblast differentiation was dependent upon HA and CD44[43]. HA production by HA synthase-2 facilitated TGF-β1-dependent fibroblast differentiation via promoting CD44 interaction with epidermal growth factor receptor (EGFR) within membrane-bound lipid rafts[43]. Our results showing that pharmacological reduction of HA by PEGPH20 or genetic deletion of CD44 ameliorated renal fibrosis (i.e. decreased HA and collagen deposition) and reduced fibroblast differentiation (i.e. decreased α-SMA and TGF-β1/Smad2/3 signalling) in HF-fed mice were consistent with these previous findings[43].

In addition, CD44 has been demonstrated to be associated with the pathogenesis of crescentic glomerulonephritis[19], AKI[20], lupus nephritis[10, 21] and CsA-induced renal injury[11]. CD44 was found to be upregulated and located in dilated tubules in the kidneys of a rat model of AKI[44]. Moreover, increased CD44 in glomerular parietal epithelial cells in aged mice contributed to glomerular hypertrophy and lower podocyte density accompanied by segmental and global glomerulosclerosis[45]. In line with these, our results showed that HF-diet feeding for 16 weeks increased renal CD44 protein expression and genetic deletion of CD44 in mice ameliorated obesity-induced tubular injury and renal dysfunction, providing evidence for a crucial role of CD44 in ORG.

The biological function of RHAMM is complex, and its extracellular and intracellular functions differ markedly. RHAMM plays a vital role in inflammation, angiogenesis[32, 33] and a variety of tissue repair processes[34, 46]. RHAMM also interacts with ERK1/2 to regulate tumour metastasis[47] and is necessary for CD44-mediated skin wound healing[48]. RHAMM is also an important target for the development of therapies via HA signalling in the tumour microenvironment[49]. RHAMM leads to progressive fibrosis and is correlated with increased severity of systemic sclerosis. Function-blocking peptides against RHAMM reduced skin fibrosis (dermal thickness and collagen production, deposition and organization), profibrotic gene expression (Tgfb1, c-Myc, Col1a1, Col3a1) and also increased the expression of the antifibrotic adipokines perilipin and adiponectin as well as prevented TGF-β1-induced myofibroblast differentiation in a bleomycin-induced mouse model of systemic sclerosis[50]. TGF-β1 was found to stimulate RHAMM expression[51]. However, the role of RHAMM in ORG has not been studied. In the present study, we observed that HF-diet feeding in mice increased RHAMM protein expression and global *Hmmr* deletion in HF-fed mice significantly reduced renal fibrosis, decreased glomerular areas, improved tubular injury and renal dysfunction. Our data demonstrated a vital role of RHAMM in ORG for the first time.

The formation of a triple complex between HA, CD44 and RHAMM on the cell surface during tumorigenesis had been reported[52]. The role of RHAMM was to enhance CD44 surface localization, stabilize the HA-CD44 interaction and contribute to the activation of ERK1/2 signalling[53]. It has been reported that cell surface RHAMM associates with several integral protein and non-protein tyrosine kinase receptors including TGF Receptor-1[32], CD44[48] and CD44-EGFR complexes[43, 54], to regulate HA deposition and CD44 and RHAMM protein expression. In our study, HF diet-induced increases in CD44 protein expression was abolished in HF-fed *Hmmr* knockout mice. We speculate that beneficial action of *Hmmr* deletion was associated with reduced cell surface HA-CD44-RHAMM complexes, which however, remains to be further explored.

Notably, PEGPH20 treatment or genetic deletion of CD44 had no effect on the glomerular area in HF-fed mice despite improved renal fibrosis and function. However, HF diet-induced increases in glomerular area was reversed by global *Hmmr* gene deletion. PEGPH20 treatment or genetic deletion of CD44 did not affect RHAMM expression. But HF diet-induced increase in CD44 protein expression in *Hmmr* ^+/+^ mice was abolished by global *Hmmr* gene deletion. These results suggest that reversal of glomerular area in obese mice may require the reduction of both CD44 and RHAMM proteins. While CD44 expression was increased in insulin-resistant podocytes, PT cells and mesangial cells, RHAMM was only increased in insulin-resistant podocytes. Therefore, CD44 and RHAMM may have distinct and cell-specific roles in regulating kidney function.

Obesity is a chronic low-grade inflammatory condition with up-regulation of proinflammatory cytokines (e.g. TNF-α, IL-6, IL-1β, MCP-1) and free fatty acids in the circulation, and activation of inflammatory pathways (e.g. P38/JNK MAPK pathways), which contribute to kidney hypertrophy and dysfunction. Our results support this concept, where renal fibrosis and dysfunction was associated with elevated inflammation (i.e. increased expression of pro-inflammatory cytokines, decreased expression of anti-inflammatory cytokines, and decreased phosphorylation of P38/JNK), and vice versa in the mouse models that were tested in the present study. The MAPK signalling pathway is a positive regulator of genotoxic stress that can subsequently induce cell apoptosis[55]. Consistent results were reported by Amos et al., who found that inhibition of p38/JNK activation reduced kidney inflammation and fibrosis in rat crescentic glomerulonephritis[56].

There is mounting evidence that early ECM remodelling is associated with obesity-associated insulin resistance. Previous studies by us and others have shown this to be the case in skeletal muscle, liver, and adipose tissue where insulin resistance is associated with increased deposition of ECM components (e.g. collagens and HA) and their interaction with membrane bound receptors such as integrins and CD44 in obese mice[5, 57–60]. The kidney also responds to the hormone insulin and reduced insulin action in podocytes leads to impaired glomerular and renal function[61]. In addition, insulin resistance prevails in patients with CKD and contributes to the progression of renal disease[62]. The relationship between insulin resistance and CKD also extends to ORG. Patients with ORG exhibited approximately 45% podocyte loss and insulin resistance specifically in podocytes triggers podocyte morphology changes and apoptosis[63]. Here we showed that *in vitro* obesogenic and insulin-resistant human kidney cells also underwent ECM remodelling, evidenced by changes in HA catabolism, increased deposition of collagens, changes in collagen modification, as well as increased expression of membrane ECM receptors including integrins, CD44 and RHAMM. Remarkably, these remodifications are cell type specific. While HA remodelling, collagen deposition and ECM receptor upregulation primarily occurred in podocytes and PT cells, collagen modification and crosslinking occurred in GEnC cells. These results support our concept that ECM-receptor activation contributes to insulin resistance[5, 57], as podocytes and PT cells are the main cell types responsive to insulin in the kidney[64]. However, the importance and relative contribution of each cell type to the pathological changes in ORG is unknown and remains to be elucidated.

In conclusion, this is the first study to demonstrate that HA and its receptors, CD44 and RHAMM mediate obesity-induced renal inflammation, fibrosis, and dysfunction, possibly via activation of TGF-β1/Smad2/3, P38/JNK MAPK and ROCK/ERK pathways. We further provided evidence that pharmacological and genetic ablation of these molecules in mice reversed adverse renal effects of obesity, therefore an intervention targeting the HA-CD44/RHAMM pathway may represent a novel therapeutic strategy against the progression of obesity-induced kidney injury. Mechanistic work in insulin-resistant human kidney cells *in vitro* illustrated an association between renal ECM remodelling and insulin resistance. Further studies to elucidate the causal relationship between renal ECM remodelling, kidney insulin resistance, and subsequent obesity-related kidney disease are warranted. Lastly, we showed clinical relevance of CD44 and RHAMM in human CKD and diabetic nephropathy, highlighting their implications and therapeutic potential in wider human kidney diseases.

## Methods

### Animal experiments

Animal work was conducted in accordance with the United Kingdom Animals (Scientific Procedures) Act 1986, the ARRIVE guidelines and the University of Dundee Welfare and Ethical Use of Animals Committee. All Mice were housed in an air-conditioned room at 22±2°C with a 12-h light/12-h dark cycle and had free access to water and food. All mice used in these experiments were male and were fed either a high fat (HF) diet (60% calories as fat, SDS #824054) or maintained on a chow diet (13% calories as fat, DBM; #D/811004) starting at 6 weeks of age for 16 weeks. All mice were studied at 22 weeks of age. Only male mice were studied due to their robust response to HF diet-induced obesity and kidney morphological changes, therefore the current study may limit its clinical relevance to the male gender.

*Cd44-*null mice (*Cd44*^-/-^) were obtained on a C57BL/6 background from Jackson Laboratory (Maine; stock no.005085). *Hmmr*-null mice (*Hmmr*^-/-^) [RHAMM knock-out] on the C57BL/6 background were a kind gift from Dr Eva Ann Turley (London Health Sciences Centre, London, Canada; The University of Western Ontario, London, ON, Canada) and generated as previously described[65]. For the reduction of HA content, mice received injections of either vehicle (10mmol/L histidine, 130mmol/L NaCl at pH6.5) or PEGylated recombinant human hyaluronidase PH20 (PEGPH20; Provided under a Material Transfer Agreement with Halozyme Therapeutics, San Diego, CA) at 1 mg/kg through the tail vein, once every 3 days for 28 days.

Three different experimental setups were used to examine the role of HA and its receptors in obesity-associated kidney injury, where mice were divided into 4 groups respectively. **Study 1:** (1) Normal “Chow” diet-fed C57BL/6 mice (Chow); (2) HF diet-fed C57BL/6 mice (HF); (3) HF diet-fed mice receiving vehicle injections (HF Vehicle); (4) HF diet-fed mice receiving PEGPH20 injections (HF PEGPH20). **Study 2:** (1) HF diet-fed CD44 wildtype littermate control mice (HF *Cd44*^+/+^); (2) HF diet-fed *Cd44*-null mice (HF *Cd44*^-/-^); (3) HF diet-fed *Cd44*-null mice with vehicle injections (HF *Cd44*^-/-^ Vehicle); (4) HF diet-fed *Cd44*-null mice with PEGPH20 injections (HF *Cd44*^-/-^ PEGPH20). **Study 3:** (1) Chow diet-fed RHAMM wildtype littermate control mice (Chow *Hmmr* ^+/+^); (2) Chow diet-fed *Hmmr*-null mice (Chow *Hmmr* ^-/-^); (3) HF diet-fed wildtype control mice (HF *Hmmr* ^+/+^); (4) HF diet-fed *Hmmr*-null mice (HF *Hmmr* ^-/-^). At the time of sacrifice, blood and kidney tissue samples were collected. For each mouse, one kidney was snap-frozen at -80°C for gene expression and protein analysis and the other was fixed in 10% formalin for histology.

### Pressure-Volume (PV) loop analysis

Pressure volume loop is considered a gold standard for the measurement of cardiac function. Mice were anaesthetized using 2% isoflurane (volume/volume). Admittance catheter (1.2F, Transonic) coupled to ADV500 data acquisition system (Transonic) visualised by LabChart (ADInstruments) sensors, was introduced into the aorta via the carotid artery to measure arterial pressure. During the procedure body temperature of mice was monitored by a rectal thermometer probe. The blood pressure data obtained from the experimental mice were analysed using Lab Chart Pro 8 software (ADInstruments).

### Renal function measurement

Serum creatinine level was determined using a Creatinine Assay kit (#ab65340, Abcam). Creatinine concentration was calculated as: *Creatitine concentration* = (amount of Creatinine nmol × Creatinine molecular weight)/ sample volume µl × sample dilution factor.

### Histology and immunohistochemistry

Paraffin-embedded mouse kidney sections (5-μm thickness) were prepared by a routine procedure (3 min in histoclear, three times; 3 min in 100% ethanol, three times; 1 min in 95% ethanol; 1min in 80% ethanol; and 5 min in distilled water). Sections were stained with Periodic acid-Schiff (PAS) staining, using the PAS Stain Kit (#ab150680; Abcam), and Sirius red staining, using Direct Red 80 (#365548; Sigma) and Picric acid (#P6744; Sigma). Immunohistochemical staining of α-SMA was performed using anti-α-SMA(#D4K9N, Cell signaling, 1:200) as previously described[5], HRP-linked Anti-rabbit (#7074S, Cell signaling, 1:1000) and DAB Substrate kit (#D3939, Sigma). HA was assessed as previously described[5], using a biotinylated HA-binding protein (#AMS.HKD-BC41, amsbio, 1:200). All staining procedures and image analysis were carried out in a blinded manner. Images were captured using camera mounted on an AxioVision microscope (Zeiss Axioscope, Germany). Quantification of Sirius red, α-SMA and HA staining were assessed in the cortex and outer medullar area using Image J Software. For PAS staining, the glomerular areas were determined by randomly analysing 6 glomeruli in the outer cortex in each kidney section. The kidney cortex and outer medulla were examined for the tubular epithelial cell vacuolar deformation/hypertrophy, tubular dilation, loss of brush border and cell lysis. The degree of tubular damage was calculated using the scoring system described by Haut et al[66].

### RNA extraction and real time RT-PCR

Total RNA was isolated from mouse kidneys using TRIzol reagent (#20130301, Ambion) and reversely transcribed into cDNA using Superscript II 10000U (#2409783, Invitrogen). The mRNA expression levels were determined by real-time quantitative RT-PCR using an Applied Biosystems (QuantStudio 7 Flex, Thermofisher, USA). The sequences of specific primers are listed in Supplemental Table 1. Data were normalized to the 18S expression and quantified using the 2^-ΔΔCT^ method.

### Western blotting

Kidney homogenates were prepared in 1 × Lysis buffer containing 92mg/ml Sucrose, 0.1% β-Mercaptaethanol, 1mM Na_3_VO_4_, 1mM Benzamidine, and 0.1M PMSF using 0.5mm zirconium oxide beads (#E-1626, ZROB05; Next Advance). The supernatant was collected after centrifugation at 9000 rpm for 5 min. The protein concentration was measured using a Bradford Reagent (#B6916; Sigma). Protein samples (40μg) were run on a 10% SDS-page gel (at 150 V) and transferred to nitrocellulose membranes (#10600002, Lytiva Amersham^TM^). The following primary antibodies were used to detect respective proteins by incubation overnight at 4°C in 5% BSA: CD44 (#E7k2y, Cell signaling, 1:1000), RHAMM (#E7S4Y, Cell signaling, 1:1000), TGF-β1 (#ab179695, Abcam, 1:1000), Phospho-Smad2 (Ser^465/467^)/Smad3 (Ser^423/425^) (#8828S, Cell signaling, 1:1000), Smad2/3 (#3102S, Cell signaling, 1:1000), Phospho-p38 MAPK (Thr^180^/Tyr^182^) (#9211, Cell signaling, 1:1000), p38 MAPK (#9212, Cell signaling, 1:1000), Phospho-SAPK/JNK (Thr^183^/Tyr^185^) (#9251, Cell signaling, 1:1000), SAPK/JNK (#9252, Cell signaling, 1:1000), Phospho-Akt (Thr^308^) (#4056, Cell signaling, 1:1000), Akt (#9272, Cell signaling, 1:1000), Phospho-p44/42 MAPK (Erk1/2) (Thr^202^/Tyr^204^) (#4370, Cell signaling, 1:1000), p44/42 MAPK (Erk1/2) (#4695, Cell signaling, 1:1000), ROCK2 (#8236, Cell signaling, 1:1000), GAPDH (#D16H11, Cell signaling, 1:1000), and Tubulin (#ab6046, Abcam, 1:1000). The membranes were then washed six times with TBST containing 0.1% Tween-20 and incubated in either HRP-goat-anti-sheep (#XDP1015101, RD SYSTEMS) or HRP-goat-anti-rabbit (#D11103-05, LI-COR) secondary antibodies for 1 h at room temperature. Densitometric quantification was performed on scanned images using the ImageJ studio.

### Cell culture and induction of insulin resistance

Conditionally immortalized human podocytes were maintained in RPMI-1640 containing L-glutamine and NaHCO_3_, supplemented with 10% FBS (Sigma Aldrich, Gillingham, UK). Proximal tubular (PT) cells were cultured in DMEM-HAM F-12 (Lonza) containing 36ng/ml hydrocortisone (Sigma) 10ng/ml EGF (Sigma), 40pg/ml Tri-iodothyronione (Sigma) and 10% Fetal bovine Serum (FBS). Glomerular Endothelial Cells (GEnC) were maintained in Endothelial Cell Growth Medium-2, containing microvascular SingleQuots Supplement Pack in 5% FBS (Lonza). Cells were studied after 12-14 days of differentiation at 37°C and were free of *Mycoplasma* infection. To induce obesogenic and insulin resistant condition, cells were cultured in the presence of 100nmol/l insulin (Tocris, Bristol, UK), 25mmol/l glucose (Sigma), 1ng/ml TNF-α and 1ng/ml IL-6 (R&D systems, Abingdon, UK). D-Mannitol (Sigma) was used as a control for osmotic pressure[37].

### Tandem Mass Tag (TMT)-Mass Spectrometry (MS) processing and analysis

Total cell protein was extracted in RIPA lysis buffer (ThermoFisher) and aliquots of each sample were digested with trypsin (2.5µg per 100µg protein; 37°C, overnight), labelled with Tandem Mass Tag (TMT) ten plex reagents according to the manufacturer’s protocol (Thermo Fisher Scientific, Loughborough, LE11 5RG, UK). Labelled samples were pooled and 50μg was desalted using a SepPak cartridge (Waters, Milford, Massachusetts, USA). Eluate from the SepPak cartridge was evaporated to dryness and resuspended in 20 mM ammonium hydroxide, pH 10, prior to fractionation by high pH reversed-phase chromatography using an Ultimate 3000 liquid chromatography system (Thermo Fisher Scientific). The sample was loaded onto an XBridge BEH C18 Column (130Å, 3.5 µm, 2.1 mm X 150 mm, Waters, UK) and peptides eluted with an increasing gradient (0-95%) of 20 mM Ammonium Hydroxide in acetonitrile, pH 10, over 60 minutes. The resulting fractions were evaporated to dryness and resuspended in 1% formic acid prior to analysis by nano-LC MSMS using an Orbitrap Fusion Lumos mass spectrometer (Thermo Scientific). High pH RP fractions were further fractionated using an Ultimate 3000 nano-LC system and spectra were acquired using an Orbitrap Fusion Lumos mass spectrometer controlled by Xcalibur 3.0 software (Thermo Scientific) and operated in data-dependent acquisition mode using an SPS-MS3 workflow. Raw data files for the Total proteome analyses were processed and quantified using Proteome Discoverer software v2.1 (Thermo Scientific) and searched against the UniProt human database (September 2018: 152,927 entries) using the SEQUEST algorithm. The reverse database search option was enabled, and all data was filtered to satisfy false discovery rate (FDR) of 5%. The data output from the Proteome Discoverer 2.1 analysis was further handled, processed and analysed using Microsoft Office Excel, GraphPad Prism and R. Normalization and differential analysis were performed in R in the same manner as RNA-seq data.

### NephroSeq Analysis

CD44 and RHAMM gene expression was analysed by Nephroseq v5 (https://nephroseq.org/) using datasets of Nakagawa CKD Kidney[67] and Ju CKD Glomeruli and Tubulointerstitium[68]. Correlation of CD44 and RHAMM gene expression with GFR, serum creatinine level and proteinuria were analysed in datasets of Ju CKD Tubulointerstitium and Schmid Diabetes Tubulointerstitium[69]. Pearson correlations were performed for statistical analysis with *p* measuring statistical significance, *r* measuring the linear correlation between two variables and *r*^2^ measuring how close the data to the fitted regression line.

### Statistics

All data except Nephroseq analyses were expressed as mean ± SEM and analysed using Prism GraphPad Software version 9. The unpaired, two-tailed Student’s t test was used to identify statistical significance between two groups. Comparisons among multiple groups were performed using one-way ANOVA followed by Tukey’s multiple comparisons test. The data were considered statistically significant at *p* < 0.05.

## Funding

This work was supported by Diabetes UK 15/0005256 and 22/0006477 (LK), British Heart Foundation PG/18/56/33935 (LK), Natural Science Foundation of Jilin Province YDZJ202301ZYTS175 (BQ), BQ and XW were supported by China Scholarship Council. This project has received funding from the Innovative Medicines Initiative 2 Joint Undertaking (JU) under grant agreement No 115974. The JU receives support from the European Union’s Horizon 2020 research and innovation programme and EFPIA and JDRF. Any dissemination of results reflects only the author’s view; the JU is not responsible for any use that may be made of the information it contains. The Medical Research Council also funded this project (Senior Clinical Fellowship to RJMC MR/K010492/1).

## Acknowledgement

PEGPH20 is an in-kind gift from Halozyme Therapeutics, Inc under a Material Transfer Agreement.

## Duality of Interest

No potential conflicts of interest relevant to this article were reported.

## Author Contributions

BQ and LK contributed to conceptualization and experimental design, researched data, contributed to discussion and data interpretation, wrote the manuscript. VM, XW, AB, CM AL, KH, and CH researched data, contributed to discussion and data interpretation, reviewed and edited the manuscript. WJ, MB and RJMC contributed to discussion and data interpretation, reviewed and edited the manuscript. All authors approved the final version of this manuscript. The authors received no editorial assistance. LK is the guarantor of this work and, as such, had full access to all the data in the study and takes responsibility for the integrity of the data and the accuracy of the data analysis.

---

**Supplemental Table 1.**
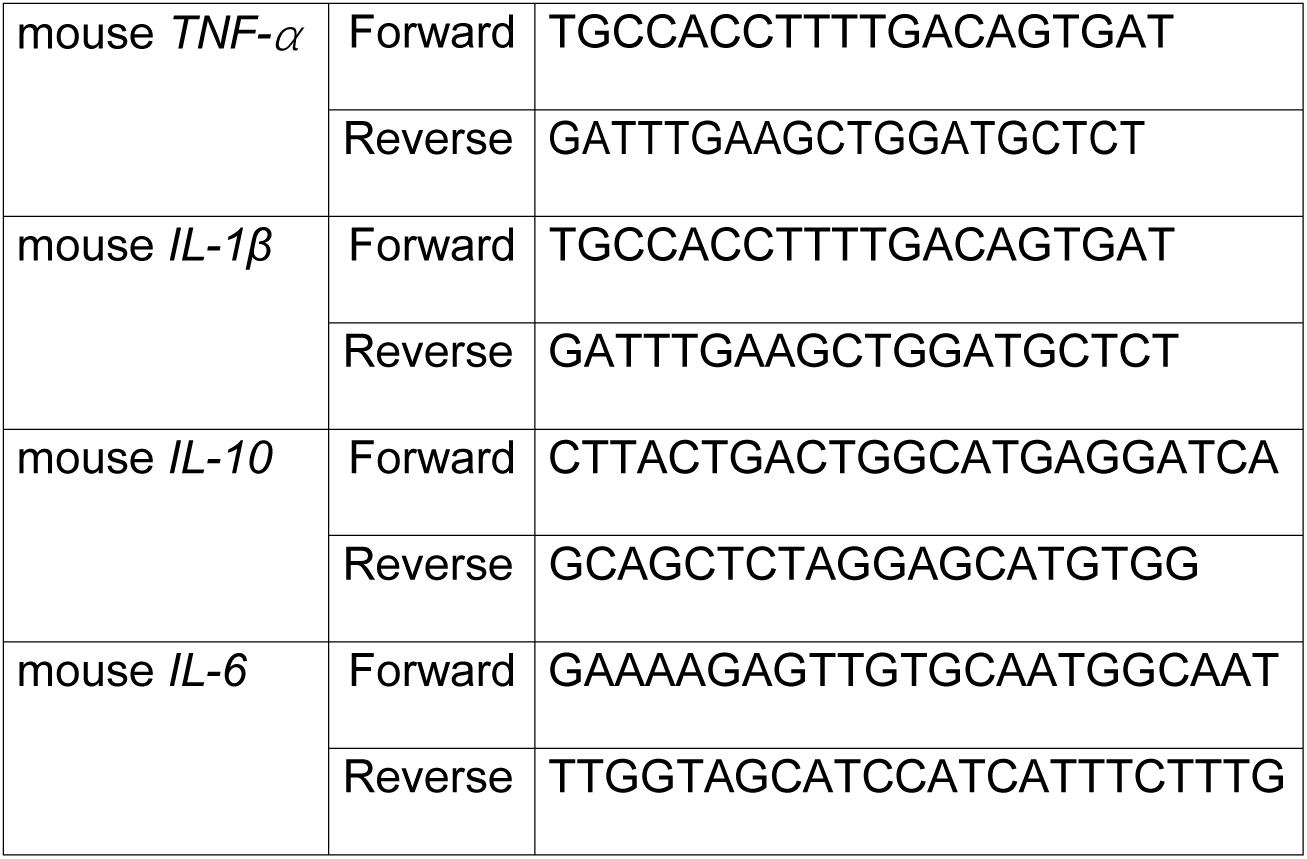
Sequences of primers used for real-time RT-PCR. *TNF-α*; tumor necrosis factor-α, *IL-1β*; interleukin 1β, *IL-10*; interleukin 10, *IL-6*; interleukin 6.

**Supplemental Fig 1.**
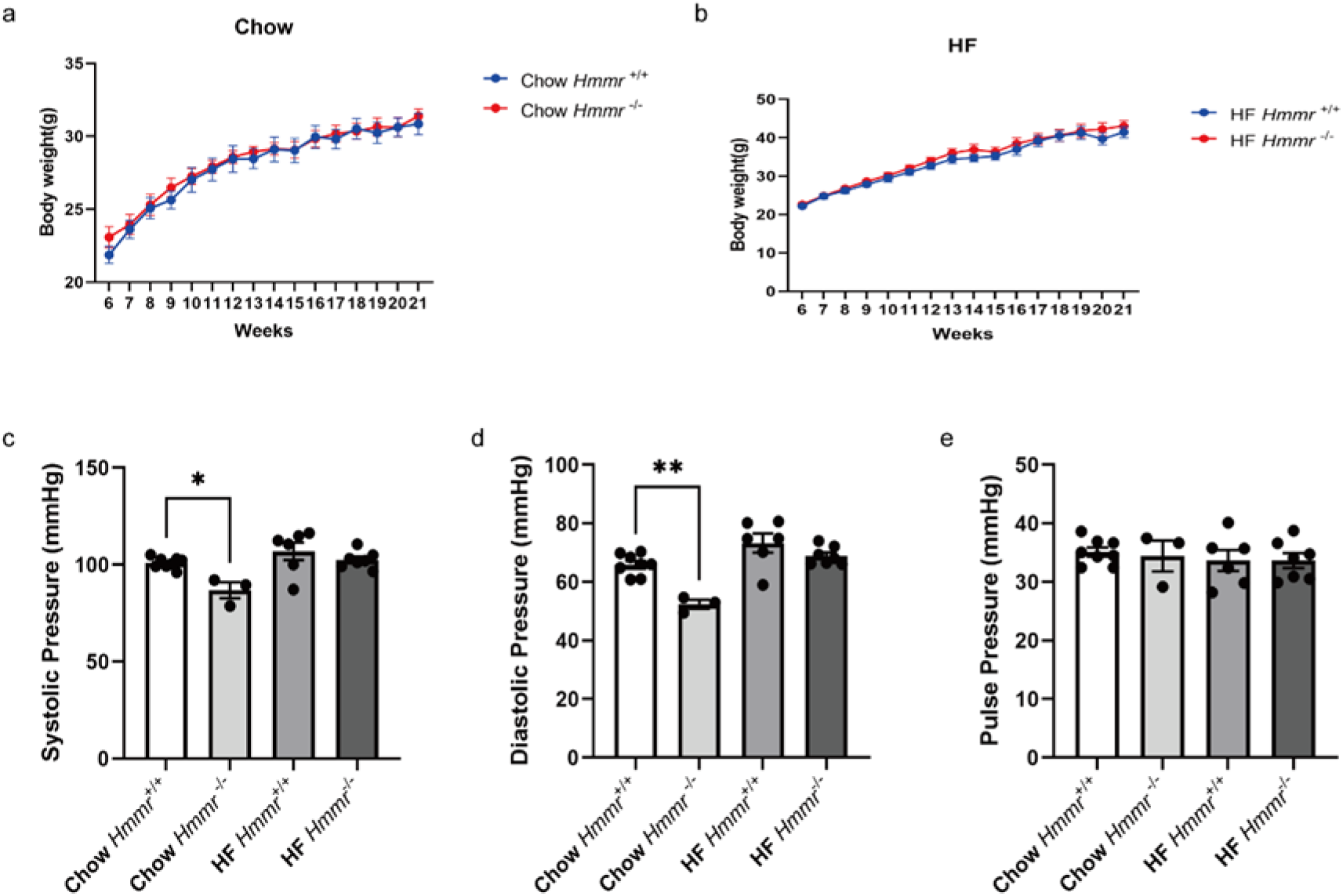
Effects of global *Hmmr* gene deletion on Body weight, Systolic, diastolic and pulse blood pressures on chow and HF diet. **a-b** Body weights of global Hmmr gene deletion mice were monitored weekly, up to 21 wk on chow and HF diets. N = 6-7 male mice. Two-way ANOVA followed by Tukey’s post-test for multiple comparisons was used for the analysis of statistical significance. **c-e** Systolic, diastolic and pulse blood pressures were monitored using a pressure volume (PV) loop, N = 3:8 for chow and N = 6:7 for HFD male mice. One-way ANOVA followed by Tukey’s post-test for multiple comparisons was used for the analysis of statistical significance. Significance \**p* < 0.05, \*\**p* < 0.01. Source data are provided in the source data file.

## Notes

### Competing Interest Statement

The authors have declared no competing interest.

## References

1. Gupta R. Epigenetic regulation and targeting of ECM for cancer therapy. Am J Physiol Cell Physiol 2022; 322(4):C762–C768.

2. Song M, Chen L, Zhang L, Li C, Coffie JW, Fang Z, et al. Cryptotanshinone enhances wound healing in type 2 diabetes with modulatory effects on inflammation, angiogenesis and extracellular matrix remodelling. Pharm Biol 2020; 58(1):845–853.

3. Jorgensen SM, Lorentzen LG, Chuang CY, Davies MJ. Peroxynitrous acid-modified extracellular matrix alters gene and protein expression in human coronary artery smooth muscle cells and induces a pro-inflammatory phenotype. Free Radic Biol Med 2022; 186:43–52.

4. Ruiz-Ojeda FJ, Mendez-Gutierrez A, Aguilera CM, Plaza-Diaz J. Extracellular Matrix Remodeling of Adipose Tissue in Obesity and Metabolic Diseases. Int J Mol Sci 2019; 20(19).

5. Hasib A, Hennayake CK, Bracy DP, Bugler-Lamb AR, Lantier L, Khan F, et al. CD44 contributes to hyaluronan-mediated insulin resistance in skeletal muscle of high-fat-fed C57BL/6 mice. Am J Physiol Endocrinol Metab 2019; 317(6):E973–E983.

6. Dogantekin E, Akgul T, Eser EP, Kotanoglu M, Bayburtluoglu V, Hucumenoglu S. The effect of intraurethral hyaluronic acid on healing and fibrosis in rats with experimentally induced urethral trauma. Int Urol Nephrol 2022; 54(4):757–761.

7. Stridh S, Palm F, Hansell P. Renal interstitial hyaluronan: functional aspects during normal and pathological conditions. Am J Physiol Regul Integr Comp Physiol 2012; 302(11):R1235–1249.

8. Wang X, Balaji S, Steen EH, Blum AJ, Li H, Chan CK, et al. High-molecular weight hyaluronan attenuates tubulointerstitial scarring in kidney injury. JCI Insight 2020; 5(12).

9. Stridh S, Palm F, Takahashi T, Ikegami-Kawai M, Hansell P. Inhibition of mTOR activity in diabetes mellitus reduces proteinuria but not renal accumulation of hyaluronan. Ups J Med Sci 2015; 120(4):233–240.

10. Yung S, Chan TM. The Role of Hyaluronan and CD44 in the Pathogenesis of Lupus Nephritis. Autoimmune Dis 2012; 2012:207190.

11. Han DH, Song HK, Lee SY, Song JH, Piao SG, Yoon HE, et al. Upregulation of hyaluronan and its binding receptors in an experimental model of chronic cyclosporine nephropathy. Nephrology (Carlton) 2010; 15(2):216–224.

12. Khunmanee S, Chun SY, Ha YS, Lee JN, Kim BS, Gao WW, et al. Improvement of IgA Nephropathy and Kidney Regeneration by Functionalized Hyaluronic Acid and Gelatin Hydrogel. Tissue Eng Regen Med 2022; 19(3):643–658.

13. Lee-Sayer SS, Dong Y, Arif AA, Olsson M, Brown KL, Johnson P. The where, when, how, and why of hyaluronan binding by immune cells. Front Immunol 2015; 6:150.

14. Heldin P, Kolliopoulos C, Lin CY, Heldin CH. Involvement of hyaluronan and CD44 in cancer and viral infections. Cell Signal 2020; 65:109427.

15. Weng X, Maxwell-Warburton S, Hasib A, Ma L, Kang L. The membrane receptor CD44: novel insights into metabolism. Trends Endocrinol Metab 2022; 33(5):318–332.

16. VerHague M, Albright J, Barron K, Kim M, Bennett BJ. Obesogenic and diabetic effects of CD44 in mice are sexually dimorphic and dependent on genetic background. Biol Sex Differ 2022; 13(1):14.

17. Decleves AE, Caron N, Nonclercq D, Legrand A, Toubeau G, Kramp R, et al. Dynamics of hyaluronan, CD44, and inflammatory cells in the rat kidney after ischemia/reperfusion injury. Int J Mol Med 2006; 18(1):83–94.

18. Chen S, Zhang M, Li J, Huang J, Zhou S, Hou X, et al. beta-catenin-controlled tubular cell-derived exosomes play a key role in fibroblast activation via the OPN-CD44 axis. J Extracell Vesicles 2022; 11(3):e12203.

19. Eymael J, Sharma S, Loeven MA, Wetzels JF, Mooren F, Florquin S, et al. CD44 is required for the pathogenesis of experimental crescentic glomerulonephritis and collapsing focal segmental glomerulosclerosis. Kidney Int 2018; 93(3):626–642.

20. Chen TH, Liu CT, Cheng CY, Sue YM, Huang NJ, Chen CH. Oligosaccharides Ameliorate Acute Kidney Injury by Alleviating Cluster of Differentiation 44-Mediated Immune Responses in Renal Tubular Cells. Nutrients 2022; 14(4).

21. Kadoya H, Yu N, Schiessl IM, Riquier-Brison A, Gyarmati G, Desposito D, et al. Essential role and therapeutic targeting of the glomerular endothelial glycocalyx in lupus nephritis. JCI Insight 2020; 5(19).

22. Fitzgerald KA, Bowie AG, Skeffington BS, O’Neill LA. Ras, protein kinase C zeta, and I kappa B kinases 1 and 2 are downstream effectors of CD44 during the activation of NF-kappa B by hyaluronic acid fragments in T-24 carcinoma cells. J Immunol 2000; 164(4):2053–2063.

23. Ferrari LF, Khomula EV, Araldi D, Levine JD. CD44 Signaling Mediates High Molecular Weight Hyaluronan-Induced Antihyperalgesia. J Neurosci 2018; 38(2):308–321.

24. Rahman AA, Soto-Avellaneda A, Yong Jin H, Stojkovska I, Lai NK, Albright JE, et al. Enhanced Hyaluronan Signaling and Autophagy Dysfunction by VPS35 D620N. Neuroscience 2020; 441:33–45.

25. Yang C, Sheng Y, Shi X, Liu Y, He Y, Du Y, et al. CD44/HA signaling mediates acquired resistance to a PI3Kalpha inhibitor. Cell Death Dis 2020; 11(10):831.

26. Mascaro M, Pibuel MA, Lompardia SL, Diaz M, Zotta E, Bianconi MI, et al. Low molecular weight hyaluronan induces migration of human choriocarcinoma JEG-3 cells mediated by RHAMM as well as by PI3K and MAPK pathways. Histochem Cell Biol 2017; 148(2):173–187.

27. Hardwick C, Hoare K, Owens R, Hohn HP, Hook M, Moore D, et al. Molecular cloning of a novel hyaluronan receptor that mediates tumor cell motility. J Cell Biol 1992; 117(6):1343–1350.

28. Savani RC, Wang C, Yang B, Zhang S, Kinsella MG, Wight TN, et al. Migration of bovine aortic smooth muscle cells after wounding injury. The role of hyaluronan and RHAMM. J Clin Invest 1995; 95(3):1158–1168.

29. Jaskula K, Sacharczuk M, Gaciong Z, Skiba DS. Cardiovascular Effects Mediated by HMMR and CD44. Mediators Inflamm 2021; 2021:4977209.

30. Misra S, Hascall VC, Markwald RR, Ghatak S. Interactions between Hyaluronan and Its Receptors (CD44, RHAMM) Regulate the Activities of Inflammation and Cancer. Front Immunol 2015; 6:201.

31. Chi A, Shirodkar SP, Escudero DO, Ekwenna OO, Yates TJ, Ayyathurai R, et al. Molecular characterization of kidney cancer: association of hyaluronic acid family with histological subtypes and metastasis. Cancer 2012; 118(9):2394–2402.

32. Park D, Kim Y, Kim H, Kim K, Lee YS, Choe J, et al. Hyaluronic acid promotes angiogenesis by inducing RHAMM-TGFbeta receptor interaction via CD44-PKCdelta. Mol Cells 2012; 33(6):563–574.

33. Hauser-Kawaguchi A, Luyt LG, Turley E. Design of peptide mimetics to block pro-inflammatory functions of HA fragments. Matrix Biol 2019; 78–79:346-356.

34. Ma X, Pearce JD, Wilson DB, English WP, Edwards MS, Geary RL. Loss of the hyaluronan receptor RHAMM prevents constrictive artery wall remodeling. J Vasc Surg 2014; 59(3):804–813.

35. Kang L, Lantier L, Kennedy A, Bonner JS, Mayes WH, Bracy DP, et al. Hyaluronan accumulates with high-fat feeding and contributes to insulin resistance. Diabetes 2013; 62(6):1888–1896.

36. Balzer MS, Rohacs T, Susztak K. How Many Cell Types Are in the Kidney and What Do They Do? Annu Rev Physiol 2022; 84:507–531.

37. Lay AC, Hurcombe JA, Betin VMS, Barrington F, Rollason R, Ni L, et al. Prolonged exposure of mouse and human podocytes to insulin induces insulin resistance through lysosomal and proteasomal degradation of the insulin receptor. Diabetologia 2017; 60(11):2299–2311.

38. Kambham N, Markowitz GS, Valeri AM, Lin J, D’Agati VD. Obesity-related glomerulopathy: an emerging epidemic. Kidney Int 2001; 59(4):1498–1509.

39. de Vries AP, Ruggenenti P, Ruan XZ, Praga M, Cruzado JM, Bajema IM, et al. Fatty kidney: emerging role of ectopic lipid in obesity-related renal disease. Lancet Diabetes Endocrinol 2014; 2(5):417–426.

40. Kim DH, Chun SY, Lee E, Kim B, Yoon B, Gil H, et al. IL-10 Deficiency Aggravates Renal Inflammation, Fibrosis and Functional Failure in High-Fat Dieted Obese Mice. Tissue Eng Regen Med 2021; 18(3):399–410.

41. Bulow RD, Boor P. Extracellular Matrix in Kidney Fibrosis: More Than Just a Scaffold. J Histochem Cytochem 2019; 67(9):643–661.

42. Kaul A, Singampalli KL, Parikh UM, Yu L, Keswani SG, Wang X. Hyaluronan, a double-edged sword in kidney diseases. Pediatr Nephrol 2022; 37(4):735–744.

43. Midgley AC, Rogers M, Hallett MB, Clayton A, Bowen T, Phillips AO, et al. Transforming growth factor-beta1 (TGF-beta1)-stimulated fibroblast to myofibroblast differentiation is mediated by hyaluronan (HA)-facilitated epidermal growth factor receptor (EGFR) and CD44 co-localization in lipid rafts. J Biol Chem 2013; 288(21):14824–14838.

44. Matsushita K, Toyoda T, Yamada T, Morikawa T, Ogawa K. Specific expression of survivin, SOX9, and CD44 in renal tubules in adaptive and maladaptive repair processes after acute kidney injury in rats. J Appl Toxicol 2021; 41(4):607–617.

45. Hamatani H, Eng DG, Hiromura K, Pippin JW, Shankland SJ. CD44 impacts glomerular parietal epithelial cell changes in the aged mouse kidney. Physiol Rep 2020; 8(12):e14487.

46. Tolg C, McCarthy JB, Yazdani A, Turley EA. Hyaluronan and RHAMM in wound repair and the “cancerization” of stromal tissues. Biomed Res Int 2014; 2014:103923.

47. Gao C, Liu S, Wang Y, Cha G, Xu X. Effect of receptor for hyaluronan-mediated motility inhibition on radiosensitivity of lung adenocarcinoma A549 cells. Transl Cancer Res 2019; 8(2):410–421.

48. Tolg C, Hamilton SR, Nakrieko KA, Kooshesh F, Walton P, McCarthy JB, et al. Rhamm-/-fibroblasts are defective in CD44-mediated ERK1,2 motogenic signaling, leading to defective skin wound repair. J Cell Biol 2006; 175(6):1017–1028.

49. Carvalho AM, Soares da Costa D, Reis RL, Pashkuleva I. RHAMM expression tunes the response of breast cancer cell lines to hyaluronan. Acta Biomater 2022; 146:187–196.

50. Wu KY, Kim S, Liu VM, Sabino A, Minkhorst K, Yazdani A, et al. Function-Blocking RHAMM Peptides Attenuate Fibrosis and Promote Antifibrotic Adipokines in a Bleomycin-Induced Murine Model of Systemic Sclerosis. J Invest Dermatol 2021; 141(6):1482–1492 e1484.

51. Amara FM, Entwistle J, Kuschak TI, Turley EA, Wright JA. Transforming growth factor-beta1 stimulates multiple protein interactions at a unique cis-element in the 3’-untranslated region of the hyaluronan receptor RHAMM mRNA. J Biol Chem 1996; 271(25):15279–15284.

52. Carvalho AM, Soares da Costa D, Paulo PMR, Reis RL, Pashkuleva I. Co-localization and crosstalk between CD44 and RHAMM depend on hyaluronan presentation. Acta Biomater 2021; 119:114–124.

53. Hamilton SR, Fard SF, Paiwand FF, Tolg C, Veiseh M, Wang C, et al. The hyaluronan receptors CD44 and Rhamm (CD168) form complexes with ERK1,2 that sustain high basal motility in breast cancer cells. J Biol Chem 2007; 282(22):16667–16680.

54. Hatano H, Shigeishi H, Kudo Y, Higashikawa K, Tobiume K, Takata T, et al. RHAMM/ERK interaction induces proliferative activities of cementifying fibroma cells through a mechanism based on the CD44-EGFR. Lab Invest 2011; 91(3):379–391.

55. Sui X, Kong N, Ye L, Han W, Zhou J, Zhang Q, et al. p38 and JNK MAPK pathways control the balance of apoptosis and autophagy in response to chemotherapeutic agents. Cancer Lett 2014; 344(2):174–179.

56. Amos LA, Ma FY, Tesch GH, Liles JT, Breckenridge DG, Nikolic-Paterson DJ, et al. ASK1 inhibitor treatment suppresses p38/JNK signalling with reduced kidney inflammation and fibrosis in rat crescentic glomerulonephritis. J Cell Mol Med 2018; 22(9):4522–4533.

57. Kang L, Ayala JE, Lee-Young RS, Zhang Z, James FD, Neufer PD, et al. Diet-induced muscle insulin resistance is associated with extracellular matrix remodeling and interaction with integrin alpha2beta1 in mice. Diabetes 2011; 60(2):416–426.

58. Williams AS, Kang L, Zheng J, Grueter C, Bracy DP, James FD, et al. Integrin alpha1-null mice exhibit improved fatty liver when fed a high fat diet despite severe hepatic insulin resistance. J Biol Chem 2015; 290(10):6546–6557.

59. Weng X, Lin, Huang JTJ, Stimson RH, Wasserman DH, Kang L. Collagen 24 alpha1 Is Increased in Insulin-Resistant Skeletal Muscle and Adipose Tissue. Int J Mol Sci 2020; 21(16).

60. Bugler-Lamb AR, Hasib A, Weng X, Hennayake CK, Lin C, McCrimmon RJ, et al. Adipocyte integrin-linked kinase plays a key role in the development of diet-induced adipose insulin resistance in male mice. Mol Metab 2021; 49:101197.

61. Kang L, Mayes WH, James FD, Bracy DP, Wasserman DH. Matrix metalloproteinase 9 opposes diet-induced muscle insulin resistance in mice. Diabetologia 2014; 57(3):603–613.

62. Spoto B, Pisano A, Zoccali C. Insulin resistance in chronic kidney disease: a systematic review. Am J Physiol Renal Physiol 2016; 311(6):F1087–F1108.

63. 63. Yang S, Cao C, Deng T, Zhou Z. Obesity-Related Glomerulopathy: A Latent Change in Obesity Requiring More Attention. Kidney Blood Press Res 2020; 45(4):510–522.

64. Artunc F, Schleicher E, Weigert C, Fritsche A, Stefan N, Haring HU. The impact of insulin resistance on the kidney and vasculature. Nat Rev Nephrol 2016; 12(12):721–737.

65. Tolg C, Poon R, Fodde R, Turley EA, Alman BA. Genetic deletion of receptor for hyaluronan-mediated motility (Rhamm) attenuates the formation of aggressive fibromatosis (desmoid tumor). Oncogene 2003; 22(44):6873–6882.

66. Hauet T, Mothes D, Goujon JM, Caritez JC, Carretier M, le Moyec L, et al. Trimetazidine prevents renal injury in the isolated perfused pig kidney exposed to prolonged cold ischemia. Transplantation 1997; 64(7):1082–1086.

67. Nakagawa S, Nishihara K, Miyata H, Shinke H, Tomita E, Kajiwara M, et al. Molecular Markers of Tubulointerstitial Fibrosis and Tubular Cell Damage in Patients with Chronic Kidney Disease. PLoS One 2015; 10(8):e0136994.

68. Ju W, Nair V, Smith S, Zhu L, Shedden K, Song PXK, et al. Tissue transcriptome-driven identification of epidermal growth factor as a chronic kidney disease biomarker. Science translational medicine 2015; 7(316):316ra193.

69. Schmid H, Boucherot A, Yasuda Y, Henger A, Brunner B, Eichinger F, et al. Modular activation of nuclear factor-kappaB transcriptional programs in human diabetic nephropathy. Diabetes 2006; 55(11):2993–3003.

